# Mathematical modelling of tissue growth control by a negative feedback

**DOI:** 10.1101/2025.01.29.635481

**Authors:** B. Kazmierczak, V. Volpert

## Abstract

This study investigates the regulation of tissue growth through mathematical modeling of systemic and local feedback mechanisms. Employing reaction-diffusion equations, the models explore the dynamics of tissue growth, emphasizing endocrine signaling and inter-tissue communication. The analysis identifies critical factors influencing the emergence of spatial structures, bifurcation phenomena, the existence and stability of stationary pulse and wave solutions. It also elucidates mechanisms for achieving coordinated tissue growth. In particular, if negative feedback is sufficiently strong, their final finite size is provided by a stable pulse, otherwise they manifest unlimited growth in the form of a wave. These findings contribute to the theoretical insights into biological processes such as embryogenesis, regeneration, and tumor development, while highlighting the role of feedback systems in maintaining physiological homeostasis.

## 1 Introduction

### 1.1 Tissue growth regulation

The proportions of the human body, including the size and relative proportions of different organs, are primarily determined by a combination of genetic factors [1] and biological signaling pathways during growth, development and regeneration [2, 3]. Genes play a critical role in determining how organs develop and maintain proper proportions relative to one another. For example, genes regulate the production of growth factors, hormones, and proteins that influence cell growth, organ formation, and the overall shape of the body. Hox genes, in particular, are a set of genes that determine the basic body plan and help control the spatial arrangement of organs and tissues during embryonic development. They ensure that organs form in the correct places and proportions.

Growth factors like fibroblast growth factor (FGF), transforming growth factor (TGF), and insulin-like growth factor (IGF) regulate cell division and growth [4, 5]. They ensure that different tissues and organs grow at appropriate rates and stop growing when they reach a certain size. Hormones such as growth hormone (produced by the pituitary gland) and thyroid hormone play critical roles in controlling growth in specific organs and overall body development [7]. For example, during puberty, growth hormone and sex hormones cause rapid growth in bones and muscles. Different organs have intrinsic growth programs that regulate their size independently to a certain degree. For example, the heart develops and grows at a rate that is proportional to the overall body size to ensure proper circulation [8]. The liver has a remarkable ability to regenerate and maintain its proportional size even if a portion is removed [9]. The body uses feedback mechanisms to regulate growth. As certain tissues grow, signals are sent to stop further growth when a critical size is reached. This prevents organs from growing too large or too small in relation to the rest of the body.

Although genetics plays an important role, environmental factors such as nutrition, physical activity, and exposure to toxins can influence how the body grows and how the organs maintain their proportion [2]. For instance, poor nutrition during childhood can lead to stunted growth and smaller organ size.

In general, a complex interaction between genetic, hormonal, and environmental factors governs the proportionality between different organs of the human body. Considering genetic and environmental factors as given, in this work we will focus on feedback mechanisms and tissue cross-talk based on the exchange of signaling molecules such as hormones and growth factors.

### 1.2 Tissue cross-talk

Tissue cross-talk refers to the communication and interaction between different tissues within an organism, particularly through signaling molecules like hormones, cytokines, growth factors, and extracellular vesicles. This communication is crucial for coordinating tissue growth, repair, and maintenance in a multicellular organism. During the regulation of tissue growth, these interactions help ensure that different tissues grow in harmony and adapt to changing physiological demands [2].

There are different mechanisms of tissue cross-talk in growth regulation. In the case of paracrine signaling, cells release signaling molecules (such as growth factors) that act on nearby cells. For instance, fibroblasts in connective tissue release growth factors like fibroblast growth factor (FGF), which promote the proliferation of epithelial cells in the skin or other tissues [10]. In endocrine signaling, hormones are released into the bloodstream and affect distant tissues. An example is growth hormone (GH) produced by the pituitary gland, which stimulates growth in bones and muscles [11]. Insulin-like growth factors (IGFs), produced in response to GH, also play a key role in tissue growth regulation [12]. Extracellular vesicles such as exosomes can carry proteins, lipids, and RNA between tissues, allowing them to communicate over long distances. These vesicles can regulate cell proliferation, differentiation, and tissue repair by transferring growth-promoting or inhibitory signals [13]. The immune system is also involved in tissue growth regulation. For example, macrophages and other immune cells release cytokines that promote or inhibit cell proliferation and tissue growth, depending on the context (e.g., during injury repair or in chronic inflammation) [14].

Among key examples of tissue cross-talk in growth regulation we can cite bone-muscle cross-talk [15]. During skeletal growth, there is coordination between bone and muscle tissues. Muscle-derived growth factors, such as myokines, influence bone development, while bone-derived factors, such as osteokines, regulate muscle growth. This interaction is crucial for proper musculoskeletal development and maintaining function throughout life.

Adipose tissue (fat) and muscle also engage in cross-talk through hormones like leptin and adiponectin [16]. Leptin, secreted by fat cells, influences energy metabolism and muscle function, while muscle-derived myokines regulate fat metabolism. This interaction is important in obesity, where dysregulation can lead to abnormal tissue growth and metabolic issues.

In cancer, the cross-talk between tumor cells and surrounding stromal tissue is a key aspect of tumor growth regulation [17]. Stromal cells can secrete growth factors and remodel the ECM to support tumor expansion, while tumor cells can influence the surrounding microenvironment to suppress immune responses or promote angiogenesis.

Tissue cross-talk plays a dynamic role in coordinating the growth and adaptation of different tissues, allowing the organism to maintain a balanced physiological state. When these interactions are disrupted, it can lead to abnormal growth patterns or diseases such as cancer, fibrosis, or tissue degeneration.

### 1.3 Models of endocrine system and tissue growth control

Endocrine axes govern and regulate the secretion of hormones from different glands through a sequence of signals. They maintain homeostasis, regulate a plethora of physiological processes (e.g. growth and development, reproductive functions), correlate responses of organisms to environmental changes (e.g. adaptation to stress). Endocrine axes rely on negative feedback loops to maintain appropriate balance between levels of different target hormones thus regulating diverse physiological phenomena. We will discuss the biological mechanisms of such feedbacks in Section 5.

Mathematical models provide indispensable tools to study tissue cross-talk and related time oscillations in neuroendocrinology [18], energy homeostasis [19], stress response [20], and other body systems [6]. One of the first ODE model of growth processes based on negative feedbacks was presented in [21]. This approach is commonly used in more recent works (see, e.g., [22]). Gradient scaling models and temporal dynamics models in the regulation of drosophila wing disk are reviewed in [2, 23, 24]. Mathematical models of tissue regeneration are presented in [25].

The main difference and the novelty of the present work is that we consider growing tissue as a spatially distributed system. This approach is quite common in tumor growth models, wound healing or morphogenesis (see, e..g., [26]) but there are few spatial models of tissue growth regulation with systemic feedback. We can cite the models of infection progression with a negative feedback by the adaptive immune response [27, 28, 29]. From the point of view of modelling and analysis, the interest of such models is that they contain integral terms and represent nonlocal reaction-diffusion equations with some different properties in comparison with the classical reaction-diffusion models.

In the next section, we will develop mathematical models of tissue growth regulation with systemic feedback. We will make abstraction of specific tissues and signaling molecules in order to develop a general framework of such models. Specific tissues and their cross-talk will be studied in the other works. Section 3 is devoted to differentiation and growth of a single tissue, and Section 4 to coordinated growth of two tissues. In Section 5, we discuss different mechanism of tissue growth control in relation to the modelling results, and conclude this paper in the last section.

## 2 Tissue growth regulation models

In this section, we will derive the tissue growth models, which will be studied in the next sections. We focus on the regulation of tissue growth through endocrine signaling involving another tissue or organ of the body. We begin with growth regulation of a single tissue and continue with the mutual regulation of two growing tissues.

### 2.1 Endocrine regulation of a single tissue

The cell concentration *u*(*x, t*) in the tissue is supposed to be described by the equation

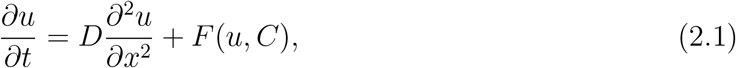

where *C* is the concentration of some biochemical substance (growth factor, hormone) regulating tissue growth. The form of the function *F* will be specified below. We take into account random cell motion described by the diffusion term.

We assume that tissue cells produce some substance with its level in the organism denoted by *B*:

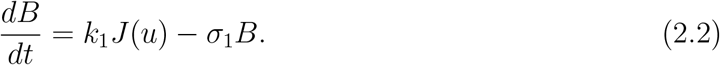

Its production rate is proportional to the total tissue volume,

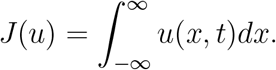

The second term in the right-hand side of equation (2.2) describes its degradation or depletion.

This substance *B* is transported to another organ or tissue by blood flow and stimulates there production of the endocrine signaling molecule *C* regulating tissue growth in equation (2.1):

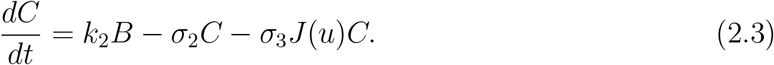

Here *C* is its concentration in the organism (or its level in blood). It acts on the cells of the tissue where its depletion is proportional to the total cell concentration *J*(*u*).

Since production and redistribution of *B* and *C* can be considered as fast compared to the cell division and death, we can use a quasi-stationary approximation in equations (2.2), (2.3) setting zero the time derivatives:

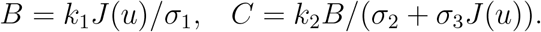

Therefore,

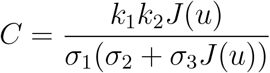

can be substituted into equation (2.1). The same reduction can be done in the analysis of the stationary solution of problem (2.1)-(2.3).

The function *F* (*u, C*) is considered in the form

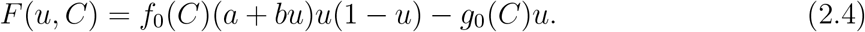

The first term in this function describes cell proliferation rate. The factor (1−*u*) represents density-dependent proliferation in which growth of cell concentration decreases their division rate. The factor (*a* + *bu*) characterizes local (autocrine, paracrine) signaling accelerating cell division. The functions *f*_0_(*C*) and *g*_0_(*C*) show the dependence of cell proliferation and death on endocrine signaling.

Substituting all these expressions in equation (2.1) we obtain the following equation

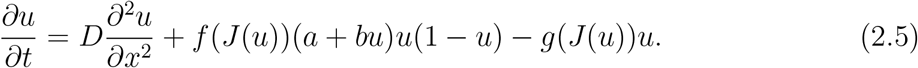

All coefficients in this equation are some positive constants. We consider it either on the whole axis or on a bounded interval with no-flux (Neumann) boundary conditions. We will specify the functions *f* (*C*) and *g*(*C*) below.

### 2.2 Interaction of two growing tissues

Consider two growing tissues with normalized concentrations of cells *u*(*x, t*) and *v*(*x, t*), respectively, and cytokines *c*_1_(*x, t*) and *c*_2_(*x, t*) produced by cells of these tissues. These concentrations are described by the following system of equations:

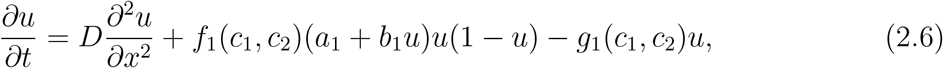

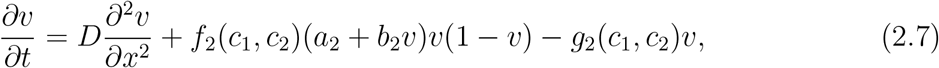

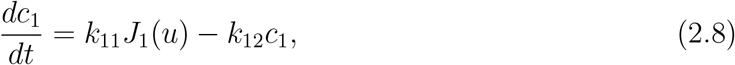

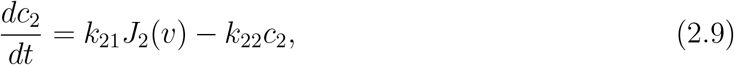

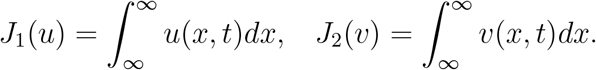

The right-hand side of equation (2.6) describes random cell motion, cell proliferation and death. Cell proliferation rate is proportional to their concentration *u* and to the logistic term (1 − *u*) describing density-dependent proliferation. The factor (*a*_1_ + *b*_1_*u*) shows that proliferation rate increases with *u* due to local cell-cell communication. Finally, the factor *f*_1_(*c*_1_, *c*_2_) shows how the proliferation rate depends on the cytokines produced by both tissues. The properties of this and other functions will be specified below. Cell death rate also depends on the concentrations of cytokines through the function *g*_1_(*c*_1_, *c*_2_). A similar equation is considered for the concentration *v*.

We suppose that all cells of the first tissue (or some proportion of all cells) produce some cytokine. We suppose that it is distributed uniformly in the organism and denote by *c*_1_ its level (quantity in a unit volume, concentration). The total quantity of this substance equals *c*_1_*V*, where *V* is the volume of the entire organism. The rate of production of this cytokine is proportional to the total cell concentration *J*_1_(*u*), its depletion or degradation is proportional to its concentration *c*_1_. The time dependence of this concentration is described by equation (2.8). A similar equation is considered for the concentration *c*_2_. We neglect here the change in total volume *V* due to tissue growth. Otherwise, the concentration *c*_1_ and the total quantity with the production rate proportional to *J*_1_(*u*) are not related by a constant factor.

Different cases according to functions *f*_*i*_, *g*_*i*_:

- Cell death is promoted by cells of the same type and down-regulated by cells of the different type, cell proliferation is independent of them,

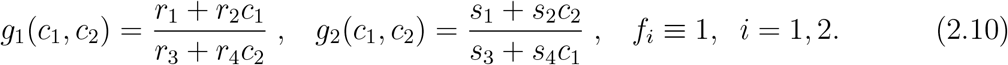
- Cell proliferation is promoted by cells of the different type and down-regulated by cells of the same type, cell death is independent of them,

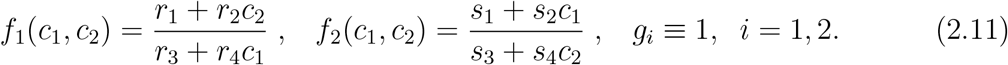
- Cell death and proliferation can depend on signaling with *c*_1_ and *c*_2_.

Summarizing these different conditions, we assume that each tissue down-regulates its own growth through a negative feedback determined by the tissue volume. On the other hand, each tissue promotes growth of the other one (Figure 2). These assumptions are confirmed by some biological observations [38]. Moreover, according to this work, cessation of developmental growth is, in the final stages, due more to an increase in the rate of cell loss than to a reduction in the rate of cell proliferation.

**Figure 1:**
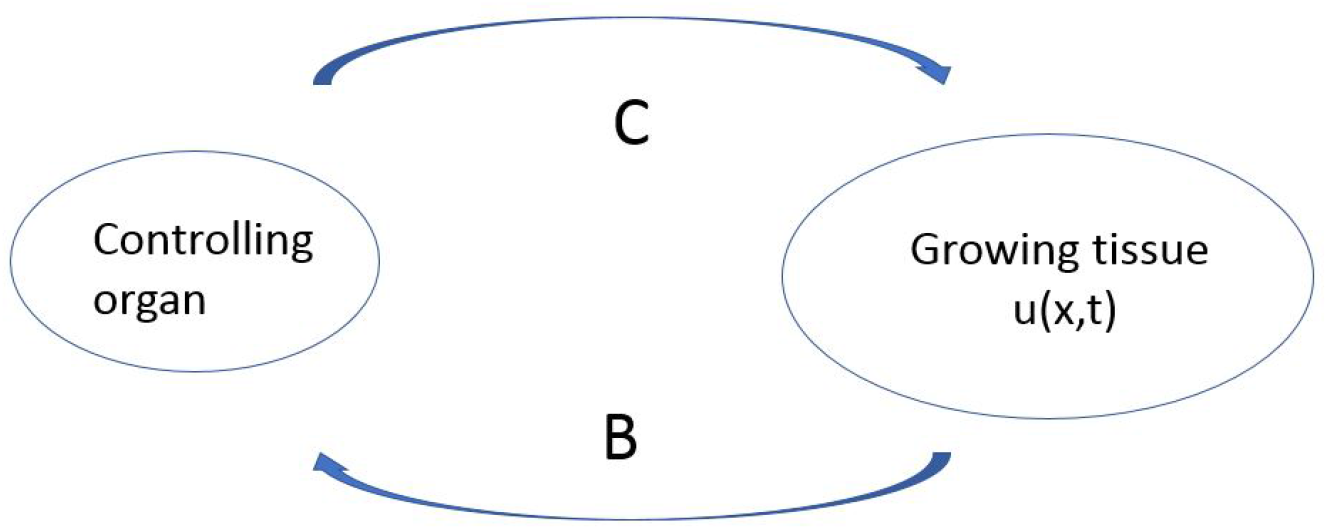
Schematic representation of the model. Growing tissue produces some signaling molecule *B* with the production rate proportional to the total cell concentration ⎰ *u*(*x, t*)*dx*. It is transported by the blood flow to the controlling organ and stimulates there production of a feedback signaling molecule *C* which can amplify or inhibit tissue growth.

**Figure 2:**
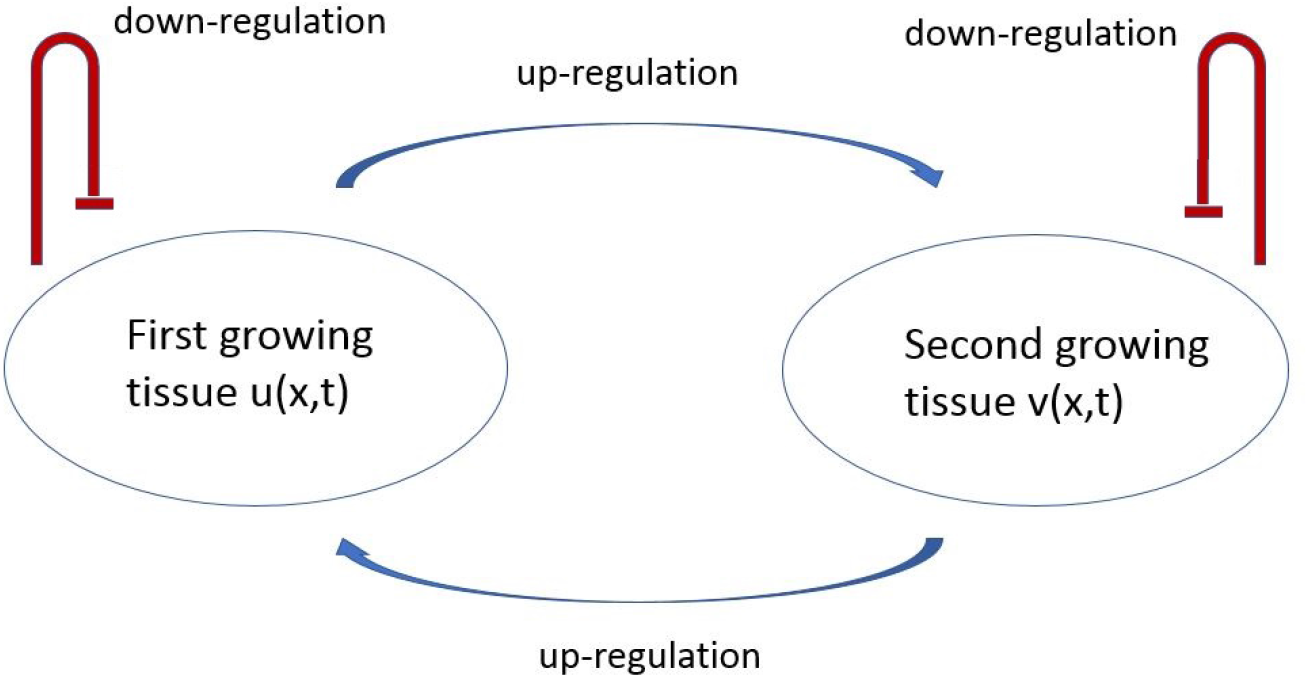
Schematic representation of the model for the coordinated growth of two tissues. Each of them down-regulates its won growth and up-regulates growth of the other one.

## 3 Differentiation and growth of a single tissue

We consider the following equation for the concentration of tissue cells *u*(*x, t*):

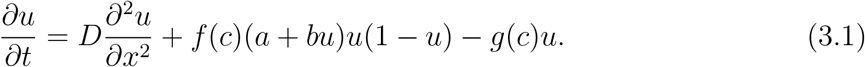

The diffusion term in the right-hand side of this equation describes random cell motion, the next term characterizes cell proliferation and the last term their death. The cell proliferation rate is considered in logistic form taking into account the decrease and arrest of the proliferation rate for the dimensionless cell concentration *u* = 1. On the other hand, cell proliferation increase due local cell-cell communication and paracrine signaling is described by the factor (*a* + *bu*). Finally, endocrine signaling on cell proliferation is described by the function *f* (*c*). Its influence on cell death is taken into account through the function *g*(*c*). These functions will be specified below. We recall that 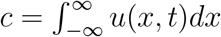 for the problem on the whole axis. For the problem on a bounded interval, the integral is taken with respect to this interval.

### 3.1. Bifurcation of spatially distributed solutions

For simplicity of calculations, we suppose that *f* (*c*) ≡ 1. Let *g*(*c*) be a non-negative continuous function defined for *c ≥* 0. The particular case *g*(*c*) = *c* is studied in [30].

Consider equation (3.1) on a bounded interval [0, *L*] with the Neumann boundary conditions:

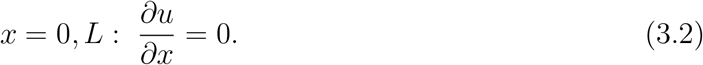

For a positive homogeneous in space stationary solution of problem (3.1), (3.2), *J*(*u*) = *Lu*, and it can be found as a solution of the equation

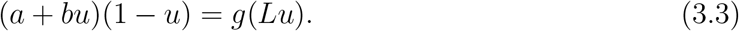

Suppose that such solution exists and denote it by *u*_0_.

We linearize equation (3.1) about this solution and obtain the eigenvalue problem:

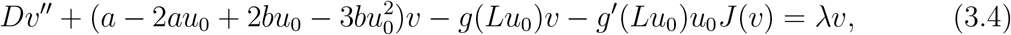

or, taking into account (3.3),

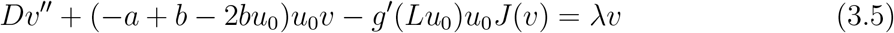

with the boundary conditions

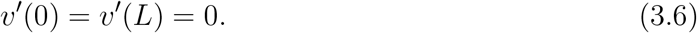

We consider the eigenfunctions

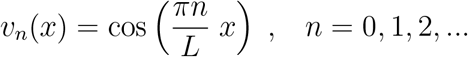

and determine the corresponding eigenvalues:

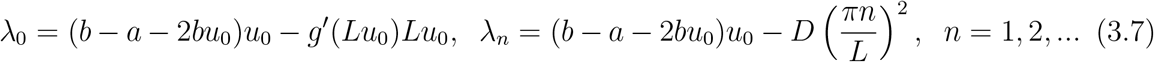

The properties of the eigenvalues are formulated in the following theorem.

#### Theorem 3.1.

*Suppose that*

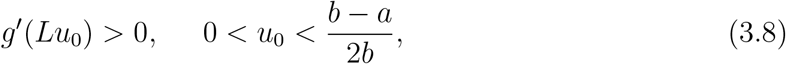

*and*

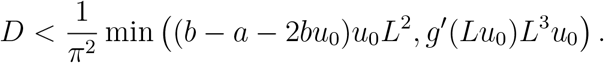

*Then λ*_1_ *is a positive eigenvalue with the maximal real part*,

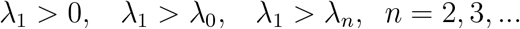

The assertion of the theorem means that the loss of stability of the homogeneous in space solution occurs with a space-dependent eigenfunction (see, e.g., [34], p. 528). Therefore, this instability leads to the bifurcation of a space-dependent solution. This is different for the equation without the integral term. In this case, *λ*_0_ *> λ*_1_, and spatial structures do not emerge (see more details in [30]).

Note that if *b > a* and *g*(*u*) = *kg*_0_(*u*), where *g*_0_(*u*) is a positive growing function such that *g*_0_(0) = 0, then the conditions of the theorem are satisfied for all *k* sufficiently large and all *D* sufficiently small.

**Example**. Let *g*(*J*) = *kJ* and *a* = 0. Then the instability conditions can be written in the following form [30]: *b/*(2*k*) *< L < b/k, D < L*^2^(2*kL − b*)(*b − kL*)*/*(*bπ*^2^). Note that the instability emerges for the values of *L* in some bounded interval. This is different in comparison with the Turing instability for which *L* should only exceed some minimal value.

Some examples of solutions bifurcating from the spatially uniform solution are shown in Figure 3. Let us note that for an asymmetric initial condition, solution converges to the half-pulse solution, while for a symmetric initial condition to the whole pulse solution. Since the initial condition is symmetric, it is orthogonal to the first eigenfunction, and the solution remains in this subspace. Since the second eigenvalue *λ*_2_ is also positive for these values of parameters, the solution converges to a symmetric pulse solution.

**Figure 3:**
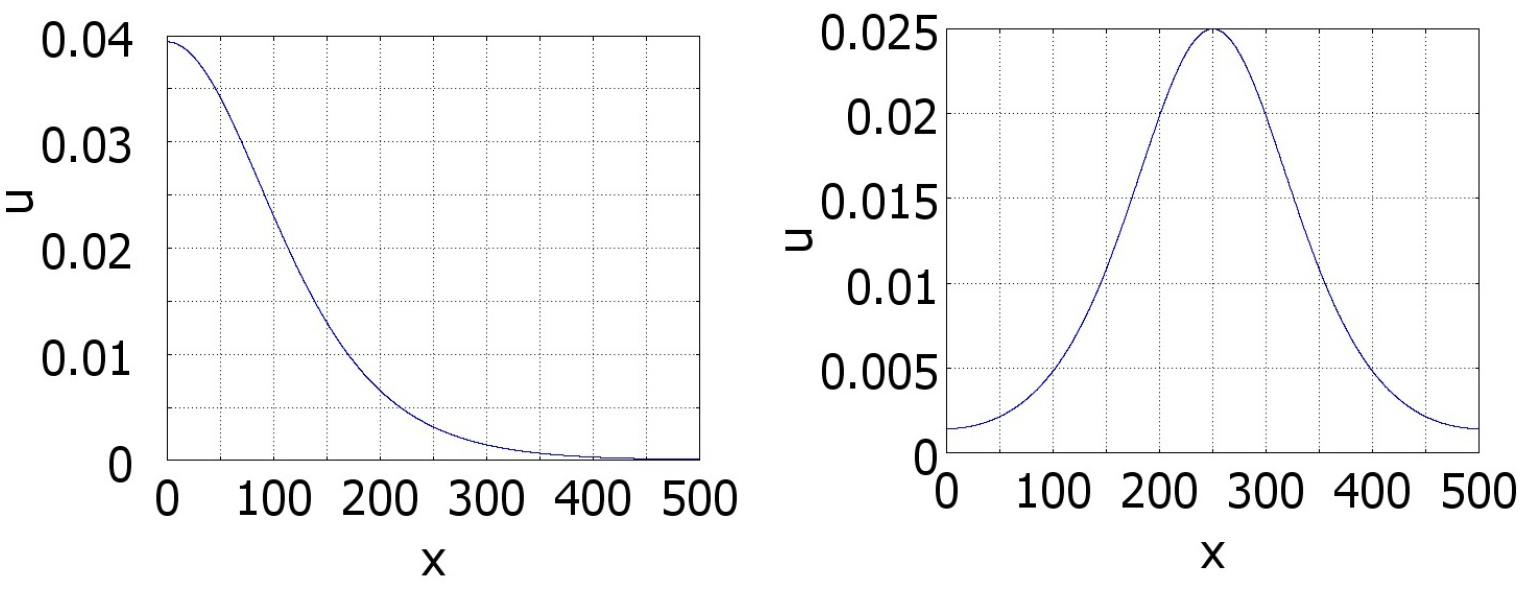
Numerical simulations of equation (3.1) with *f* (*J*) ≡ 1 and *g*(*J*) = *kJ, a* = 0.5, *b* = 1, *k* = 0.1. Left: *D* = 50, the initial data are: 0.1*H*(*x*−250 −50)*H*(250 −*x*). Right: *D* = 20, the initial data are: 0.1*H*(*x* − 250 − 25)*H*(250 + 25 − *x*).

### 3.2 Existence and stability of pulses on the whole axis

We consider the stationary equation

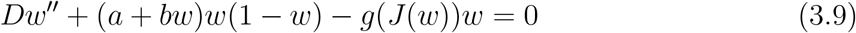

on the whole axis. We look for a positive solution of this equation vanishing at infinity,

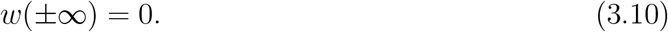

Consider the auxiliary problem

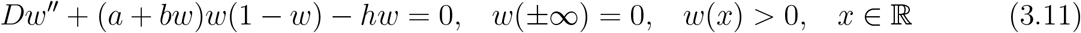

and denote by *w*_*h*_(*x*) its positive solution. Then solution of the equation

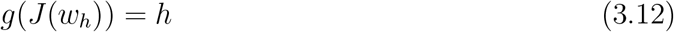

provides a solution of problem (3.9), (3.10).

Problem (3.11) has a solution if *b > a >* 0 and *a < h < h*_∗_, where *h*_∗_ *> a* is a positive number which can be determined analytically [30]. Moreover,

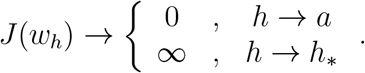

We can now formulate the existence result.

#### Theorem 3.2.

*Suppose that b > a >* 0, *g*(0) *< a, g*(∞) *> h*∗ *or g*(0) *> a, g*(∞) *< h*∗. *Then problem (3*.*9), (3*.*10) has a positive solution*.

Consider some examples.

**Examples**. If *g*(*c*) = *kc*^*n*^ with some positive *k* and *n*, then the conditions of the theorem are satisfied. They are also satisfied for negative *n*, but these two cases are different. Introduce the function *F* (*h*) = *g*(*J*(*w*_*h*_)) − *h* and consider the equation *F* (*h*) = 0. If it has a single solution *h*_0_, then *F* ^′^(*h*_0_) ≥ 0 for *n >* 0 and *F* ^′^(*h*_0_) ≤ 0 for *n <* 0. This is related to stability and bifurcations of solutions. If *n* = 0, then equation (3.9) is independent of *J*(*u*) and the pulse is unstable [31].

**Remark**. If the function *f* (*J*) is not constant, then equation

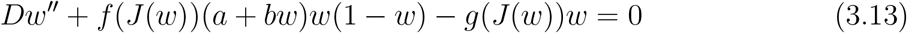

can be reduced to an equation similar to (3.9) by the introduction of a new function 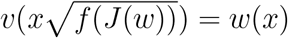.

### 3.3 Numerical simulations

Some examples of numerical simulations are shown in Figures 4 and 5. If conditions of Theorem 2.2 are satisfied, then there exists a stationary pulse solution. For the values of parameters in Figures 4 (left), it is stable, and the solution of equation (3.1) converges to it. On the contrary, for the values of parameters in Figures 4 (right), the pulse does not exist, and we observe wave propagation.

**Figure 4:**
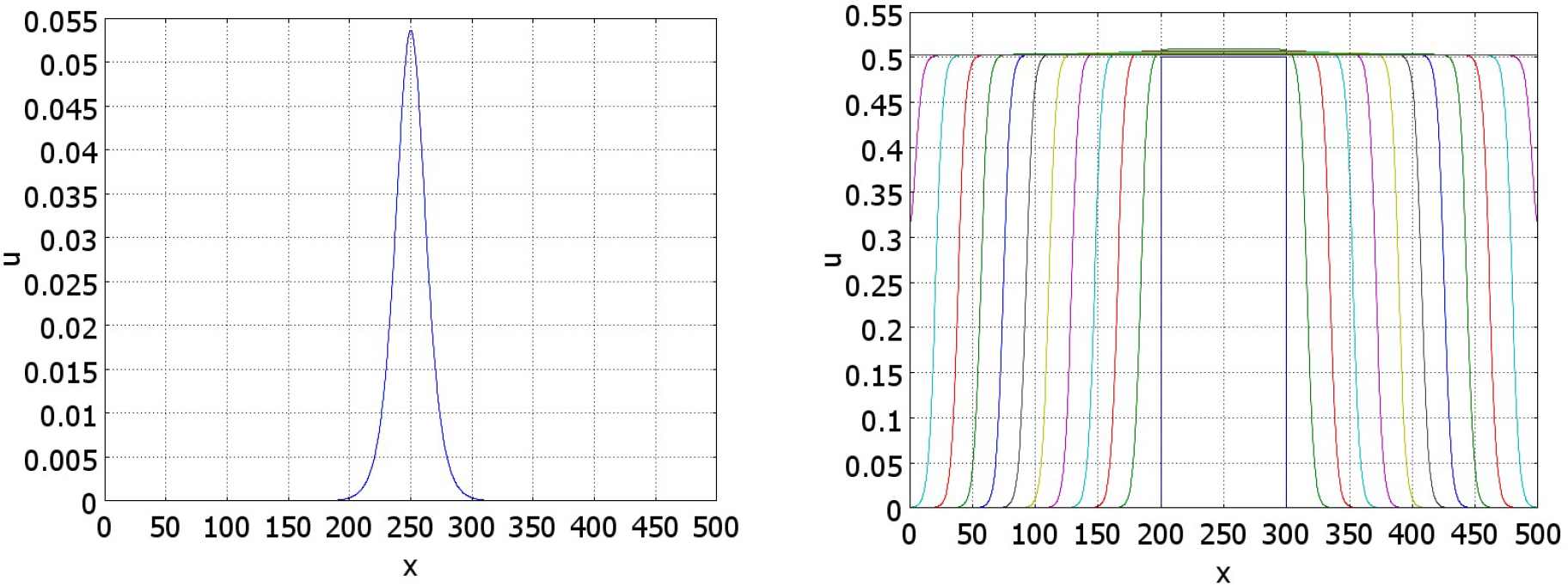
Numerical simulations of equation (3.1) with *f* (*J*) ≡ 1 and *g*(*J*) = *k*_1_ +*k*_2_*J/*(1+*J*), *a* = 0.5, *b* = 1, Left: solution converges to a stationary pulse, *k*_1_ = 0.2, *k*_2_ = 0.5, *J* = 1.7237, *t* = 1200. Right: solution propagates as a wave, *k*_1_ = 0.2, *k*_2_ = 0.1, *t* = 0, 10, …, 50.

**Figure 5:**
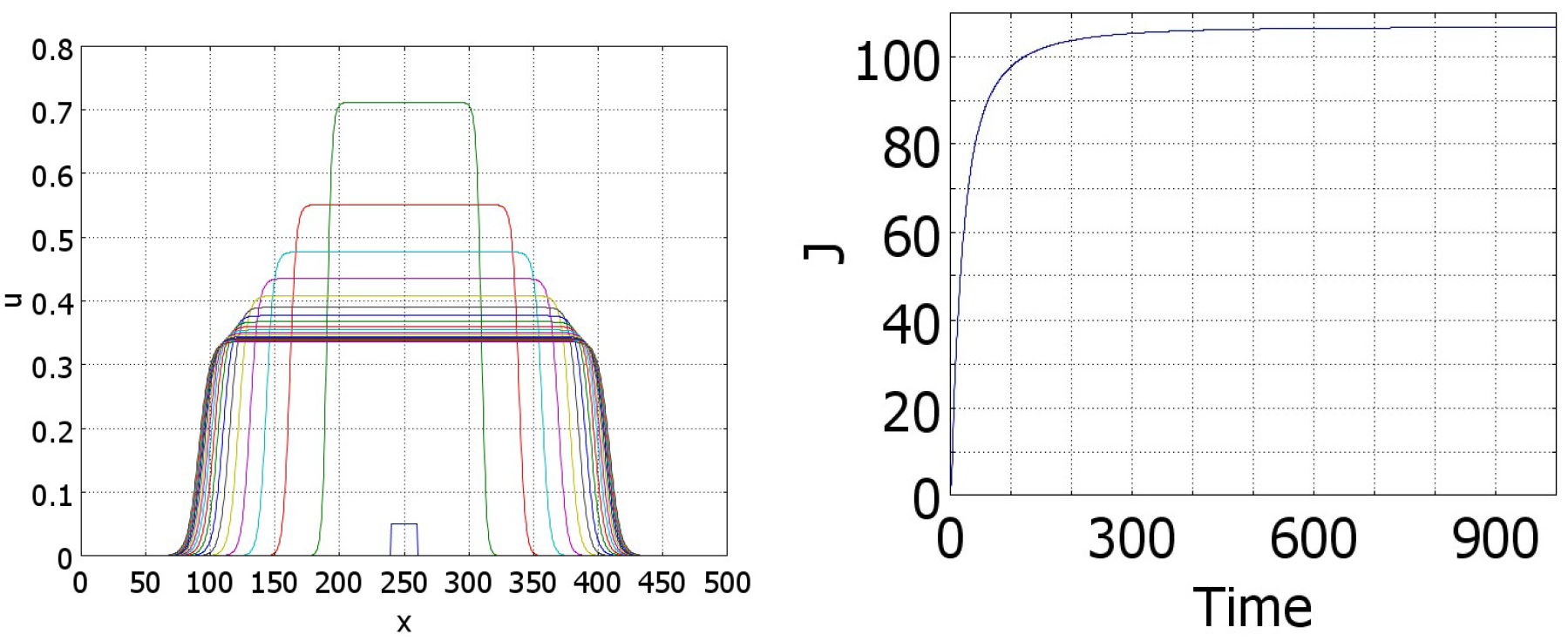
Numerical simulations of equation (3.1) with *f* (*J*) ≡ 1 and *g*(*J*) = (*k*_1_ + *k*_2_*exp*(0.05*J*)*/*(100 + *exp*(0.05*J*))), *a* = 0.5, *b* = 1, *k*_1_ = 0.05, *k*_2_ = 0.75. The left panel shows convergence of solution to the stationary pulse, *t* = 0, 50, 100, …, 1000. The corresponding function *J*(*u*)(*t*) is shown in the right panel.

Under the conditions of the theorem, if solution *h* of equation (3.12) is sufficiently close to the value *h*_∗_, then the pulse solution exists, it is wide and top-flat (Figure 5, left). The figure shows convergence of solution of the initial boundary value problem to the stationary pulse solution. Let us note that the initial condition is small. Contrary to the conventional bistable reaction-diffusion equation, here the solution can grow even for any small initial condition. In fact, the presence of the integral in the equation changes its type from the monostable case for small *J*(*u*) to the bistable case for large values. Thus, solution grows, takes the form of a flat pulse, then decreases its height and increases its width. The right panel in this figure shows the convergence of *J*(*u*)(*t*) to its limiting value.

The decrease of the plateau value in Figure 5 is determined by the density-dependent cell proliferation. If the decrease of the proliferation rate is faster, then the decrease of the plateau value is not so essential. For example, if we impose the condition that cells stop proliferation if they touch each other and proliferate otherwise, then for a uniform cell distribution, proliferation rate becomes (*a* + *bu*)*uϕ*(*u*), where *ϕ*(*u*) = *ϕ*_0_ *>* 0 for 0 ≤ *u <* 1 and *ϕ*(1) = 0. An example of such simulation is presented in Figure 6.

**Figure 6:**
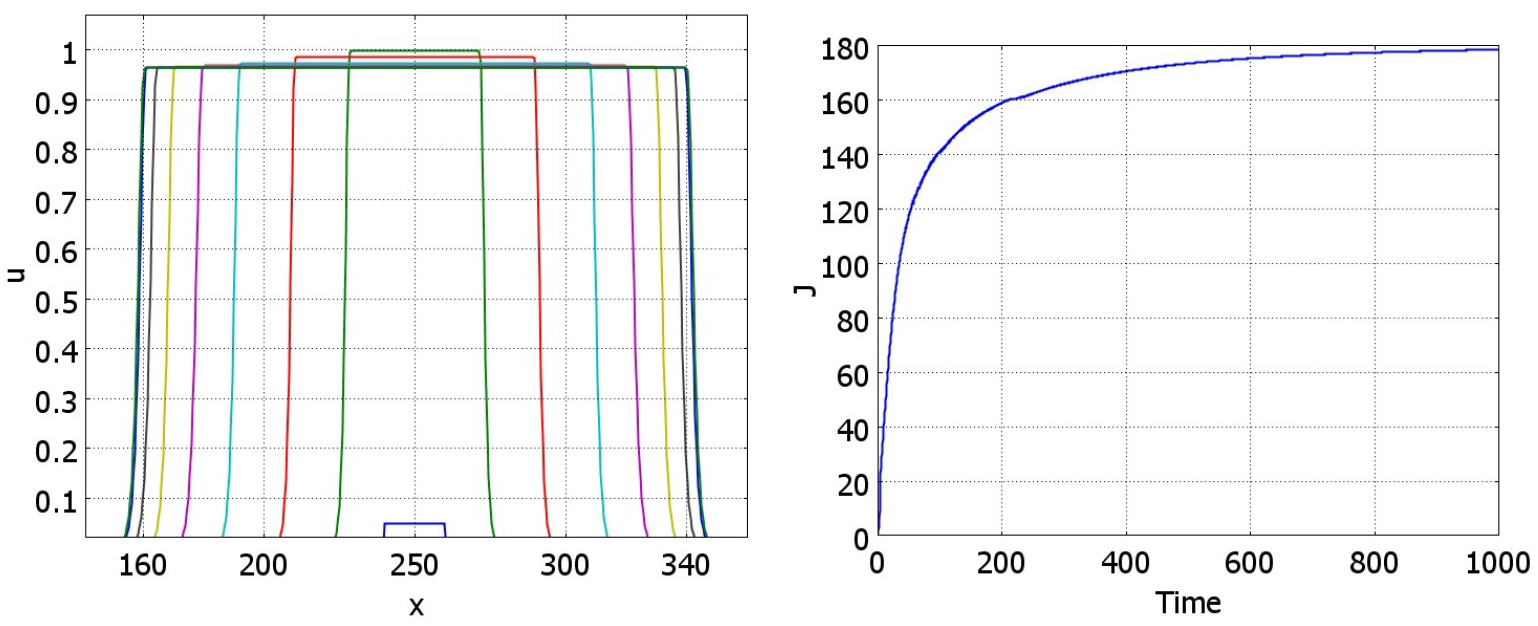
Numerical simulations of equation (3.1) with *f* (*J*) ≡ 1 and *g*(*J*) = (*k*_1_ + *k*_2_*exp*(0.05*J*)*/*(100 + *exp*(0.05*J*))), *a* = 0.5, *b* = 1, *k*_1_ = 0.05, *k*_2_ = 1.1. The left panel shows convergence of solution to the stationary pulse, *t* = 0, 12.5, 25, 50, 100, 200, 400, 8000, 1000. The corresponding function *J*(*u*)(*t*) is shown in the right panel. The proliferation rate is given by the function (*a* + *bu*)*u*(1 − *u*^100^*/*(*u*^100^ + 0.1)) with the density dependence approximating a step-wise constant function *ϕ*(*u*).

## 4 Coordinated growth of two tissues

Growth of different tissues during the development of the organism is precisely coordinated. In this section, we study simultaneous growth of two tissues which exchange signals and influence their respective growth rates and final sizes.

### 4.1 Existence of pulses

We consider quasi-stationary approximations in equations (2.8), (2.9). Then

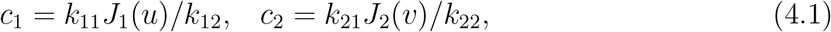

and equations (2.6), (2.7) become as follows:

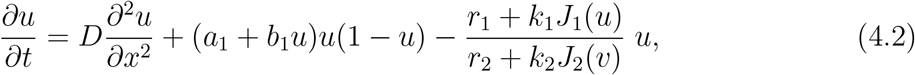

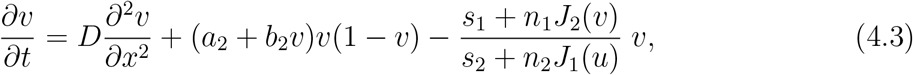

where

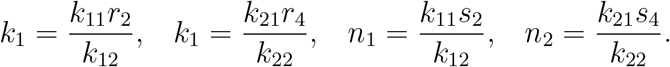

We look for a positive stationary solution *w*(*x*), *z*(*x*) of this system of equations on the whole axis:

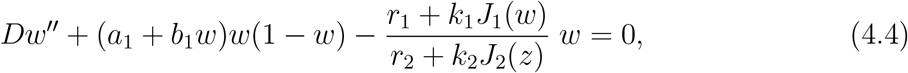

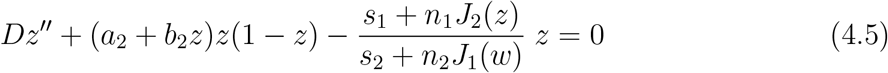

vanishing at infinity:

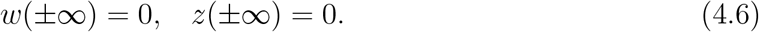

Consider the auxiliary problem

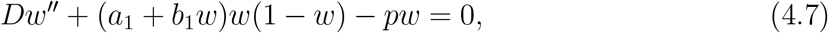

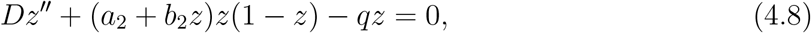

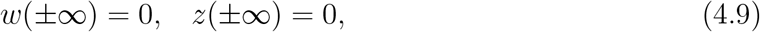

and denote its solution by (*w*_*p*_(*x*), *z*_*q*_(*x*)). This solution provides a solution of problem (4.4)-(4.6) if

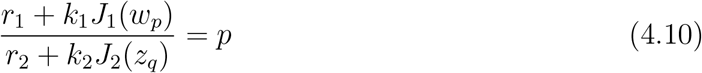

and

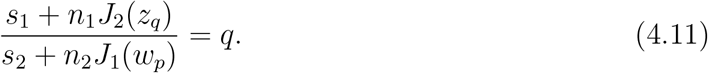

#### Lemma 4.1.

*The function J*_1_(*w*_*p*_) *is positive and continuous in the interval a*_1_ *< p < p*_∗_ *for some p*_∗_ *> a*_1_, *J*_1_(*w*_*p*_) = 0 *for p* = *a*_1_, *and J*_1_(*w*_*p*_) → ∞ *as p* → *p*_∗_. *The function J*_2_(*z*_*q*_) *is positive and continuous in the interval a*_2_ *< q < q*_∗_ *for some q*_∗_ *> a*_2_, *J*_2_(*z*_*q*_) = 0 *for q* = *a*_2_, *and J*_2_(*z*_*q*_) → ∞ *as q* → *q*_∗_.

**Proof**. By definition, *w*_*p*_(*x*) is a positive solution of equation (4.7) vanishing at infinity. It is independent of *q*. The function *F*_*h*_(*w*) = (*a* + *bw*)*w*(1 − *w*) − *hw* can have from one to three non-negative zeros. Suppose that there are three of them. This is the case if

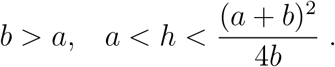

Denote by *w*_0_(*h*) the maximal solution of the equation *F*_*h*_(*w*) = 0. It is known that equation (4.7) has a positive solution *w*_*h*_(*x*) vanishing at infinity if and only if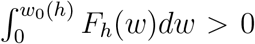. This inequality holds for *a < h < h*_∗_ = (2(*a* + *b*)^2^ + *ab*)*/*(9*b*).

Denote *w*_*m*_(*h*) = max_*x*∈ℝ_ *w*_*h*_(*x*). Then *w*_*m*_(*h*) *< w*_0_(*h*). If *h* ↗ *h*_∗_, then *w*_*m*_(*h*) ↗ *w*_0_(*h*), *w*_*h*_(*x*) → *w*_0_(*h*) uniformly in *x* on every bounded interval and, consequently, *J*_1_(*w*_*h*_) →∞ (see [30] for more detail).

Next, suppose that *h↘ a* and set *h* = *a* + *ϵ*. Then equation (4.7) can be written as follows:

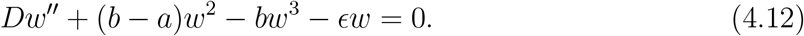

Let us introduce a new function *u*(*y*) by the equality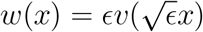. Then it satisfies the equation

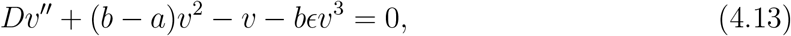

where prime denotes the derivative with respect to *y*. If *b > a* and *ϵ* is sufficiently small, then this equation has a positive solution *v*_*ϵ*_(*y*) vanishing at infinity. Then

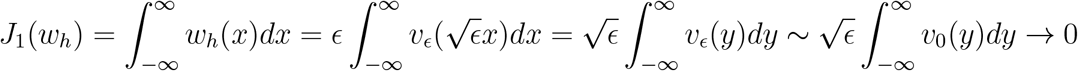

as ϵ → 0.

The second part of the lemma for *J*_2_(*w*_*q*_) can be proved similarly. The lemma is proved.

#### Lemma 4.2.

*Suppose that system (4*.*10), (4*.*11) has a solution for a*_1_ *< p < p*_∗_, *a*_2_ *< q < q*_∗_ *in some parameter range. Then J*_1_(*w*_*p*_) → ∞ *if and only if J*_2_(*z*_*q*_) → ∞.

**Proof**. Suppose that *J*_1_(*w*_*p*_) → ∞ but *J*_2_(*z*_*q*_) remains bounded. Then the left-hand side of equality (4.10) tends to infinity, while the right-hand side remain bounded since *a*_1_ *< p < p*_∗_. The second case is proved similarly. The lemma is proved.

#### Lemma 4.3.

If

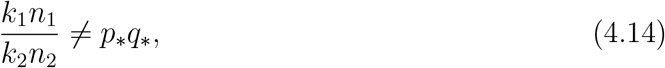

then for *p* and *q* satisfying (4.10), (4.11), convergence *J*_1_(*w*_*p*_) →∞, *J*_2_(*z*_*q*_) →∞ does not hold.

**Proof**. If this convergence occurs, then *p → p*_∗_, *q → q*_∗_. From equations (4.10), (4.11) we obtain

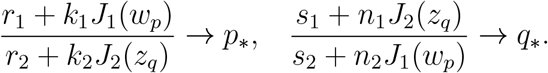

Multiplying these equations, we get

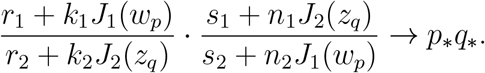

The expression in the left-hand side converges to *k*_1_*n*_1_*/*(*k*_2_*n*_2_). This contradiction proves the lemma.

We begin the analysis of the existence of solutions of system of (4.10), (4.11) with the case *r*_1_ = *r*_2_ = *s*_1_ = *s*_2_ = 0. Then, multiplying these equations, we obtain *pq* = *κ*, where *κ* = *k*_1_*n*_1_*/*(*k*_2_*n*_2_). The function *h*_2_(*p*) = *J*_2_(*z*_*κ/p*_) is defined on the interval *a*_2_ *< κ/p < q*_∗_ or *κ/q*_∗_ *< p < κ/a*_2_. Together with the function *h*_1_(*p*) = *J*_1_(*w*_*p*_), they are defined on the intersection of the intervals

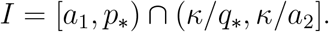

We recall that *h*_1_(*a*_1_) = 0, *h*_1_(*p*_∗_) = ∞, *h*_2_(*κ/q*_∗_) = ∞, *h*_2_(*κ/a*_2_) = 0, and these functions are positive inside their intervals of definition.

Assume that *κ/q*_∗_ *< p*_∗_ and consider the function

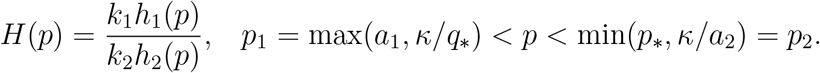

Since *h*_1_(*a*_1_) = 0 and *h*_2_(*κ/q*_∗_) = ∞, then *H*(*p*_1_) = 0. Next, equalities *h*_1_(*p*_∗_) = ∞ and *h*_2_(*κ/a*_2_) = 0 implies that *H*(*p*_2_) = ∞. Hence, equation *H*(*p*) = *p* (equivalent to equation (4.10)) has a solution in this interval. We proved the following result.

#### Proposition 4.4.

*Suppose that r*_1_ = *r*_2_ = *s*_1_ = *s*_2_ = 0 *and*

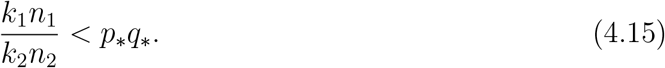

*Then problem (4*.*4)-(4*.*6) has a positive solution. If the inequality is opposite, such solution does not exist*.

We will now consider the case without the assumption that some coefficients vanish.

#### Theorem 4.5.

*If r*_1_*/r*_2_ *< a*_1_, *s*_1_*/s*_2_ *< a*_2_ *and*

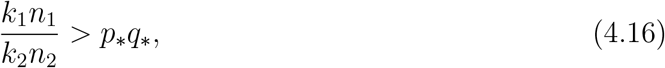

*then problem (4*.*4)-(4*.*6) has a positive solution*.

**Proof**. If *r*_1_*/r*_2_ *< a*_1_, then equation (4.10) with respect to *p* has a solution in the interval (*a*_1_, *p*_∗_) for any *q* ∈ [*a*_2_, *q*_∗_) fixed. Indeed, for *p* = *a*_1_, *J*_1_(*w*_*p*_) = 0, *J*_2_(*w*_*q*_) ≥ 0, and

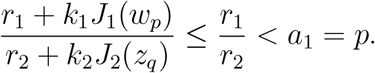

On the other hand, *J*_1_(*w*_*p*_) → ∞ as *p* → *p*_∗_, and the last inequality becomes opposite for *p* sufficiently close to *p*_∗_. Therefore, equation (4.10) has a solution for some intermediate value of *p*. Denote it by *S*(*q*). Then, *S*(*a*_2_) *> a*_1_, *S*(*q*) → *p*_∗_ as *q* → *q*_∗_.

Similarly, If *s*_1_*/s*_2_ *< a*_2_, then equation (4.11) with respect to *q* has a solution in the interval (*a*_2_, *q*_∗_) for any *p* ∈ [*a*_1_, *p*_∗_). We denote it by *T* (*p*), and *T* (*a*_1_) *> a*_2_, *T* (*p*) → *q*_∗_ as *p* → *p*_∗_.

We need to verify that the curve *p* = *S*(*q*) and *q* = *T* (*p*) intersect for some *a*_2_ *< q < q*_∗_, *a*_1_ *< p < p*_∗_. This point of intersection provides solution of system (4.10), (4.11) and, consequently, of problem (4.4)-(4.6). We will show that condition (4.16) implies that the curve *p* = *S*(*q*) is above *q* = *T* (*p*) in some vicinity of the point (*p*_∗_, *q*_∗_) (Figure 7). Then the curves intersect inside the rectangle.

**Figure 7:**
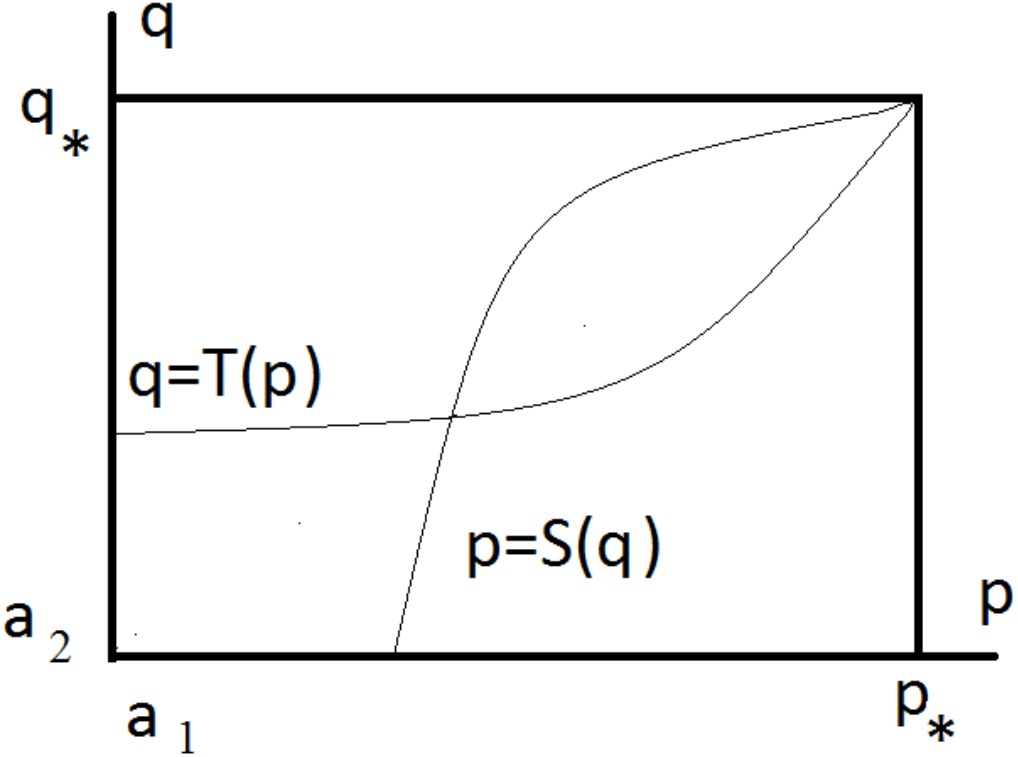
Schematic representation of the curves *p* = *S*(*q*) and *q* = *T* (*p*) on the (*p, q*)-plane. The former starts at the lower boundary of the domain and tends to the point (*p*_∗_, *q*_∗_). The latter starts at the left boundary and tends to the same point. If the first curve is above the second curve near this point, as shown in the theorem, they have a point of intersection inside the domain.

We consider the inverse function *q* = *R*(*p*) to the function *p* = *S*(*q*). For simplicity of notation, set

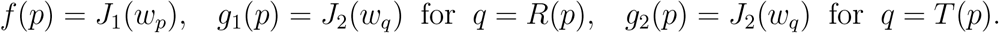

Then equations (4.10), (4.11) become as follows:

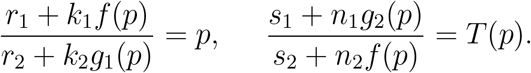

We obtain from these equations:

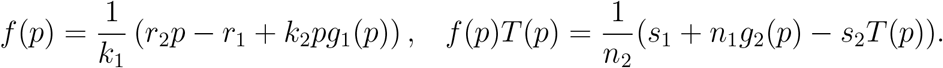

Multiplying the first equation by *T* (*p*) and equating the right-hand sides, we get the equality:

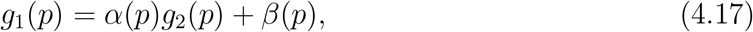

where

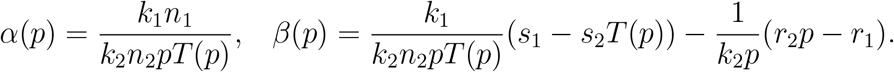

Since *p < p*_∗_, *T* (*p*) *< q*_∗_, then by virtue of condition (4.16),

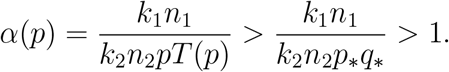

Furthermore, |*β*(*p*)| *M*, where *M* is some positive constant independent of *p*.

Equation (4.17) implies that *g*_1_(*p*) *> g*_2_(*p*) for *p* sufficiently close to *p*_∗_. Indeed, if *g*_1_(*p*_*i*_) ≤ *g*_2_(*p*_*i*_) for some sequence *p*_*i*_ such that *p*_*i*_ ↗ *p*_∗_, then from (4.17),

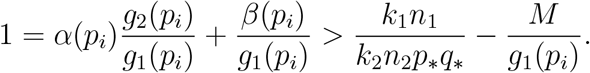

Since the first term in the right-hand side of this inequality is larger than 1, and the second one converges to 0, we obtain a contradiction.

Thus, we proved that

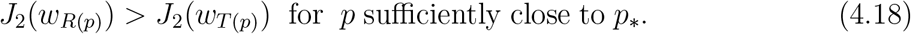

If *J*_2_(*w*_*q*_) is a monotone function of *q* for *q* sufficiently close to *q*_∗_, then we conclude that *R*(*p*) *> T* (*p*).

Monotonicity of *J*_2_(*w*_*q*_) is not needed if we want to verify that *R*(*p*_0_) *> T* (*p*_0_) for some *p*_0_. This is sufficient for the intersection of the curves and for the existence of solution. We take a value *q*_0_ of *J*_2_(*w*_*q*_) for which *J*_2_(*w*_*q*_) *<* 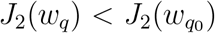if *q < q*_0_. Existence of such *q*_0_ follows from the fact that *J*_2_(*w*_*q*_) tends to infinity. Then for any *q*_1_ such that 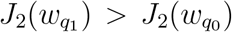 it follows that *q*_1_ *> q*_0_. We choose *p*_0_ such that *q*_0_ = *T* (*p*_0_). It follows from (4.18) that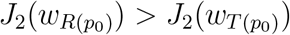. Then *R*(*p*_0_) *> T* (*p*_0_).

### 4.2 Existence of waves

System of equations (4.2), (4.3) cannot have travelling wave solutions with one of the limits at infinity different from zero since the integrals *J*_1_(*u*), *J*_2_(*v*) are not defined in this case. Instead of travelling waves in the classical definition, we will consider the solution of the Cauchy problem converging to two waves moving in the opposite directions, one to minus infinity, another one to plus infinity. In this case, the integrals are well defined. Let us proceed to the description of such solutions and to the conditions of their existence.

Consider the equations

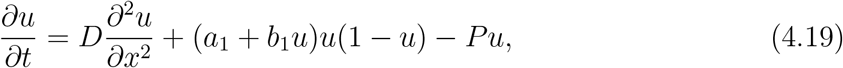

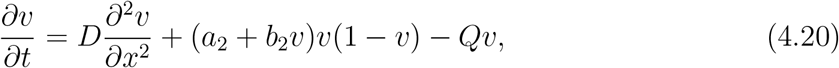

where *P* and *Q* are some positive constants. If

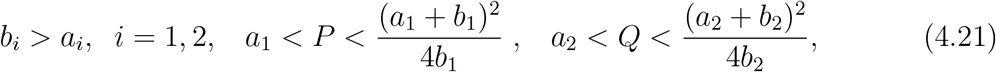

then these are bistable equations. In this case, each of the equations

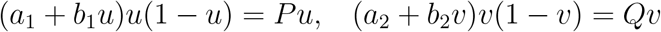

has three non-negative solutions, 0, *u*_1_, *u*_2_ and 0, *v*_1_, *v*_2_. Equations (4.19), (4.20) have travelling wave solutions with the limits [0, *u*_2_] and [0, *v*_2_]. Moreover, there are some values *p*_∗_ *> a*_1_ and *q*_∗_ *> a*_2_ (the same as in Lemma 4.1) such that the inequalities

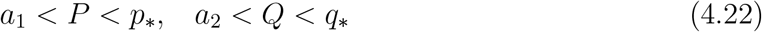

provide the positivity of the wave speeds. Denote the wave speeds by *c*_1_ and *c*_2_, respectively. If 0 *< P < a*_1_, then the corresponding equation is monostable, the wave still exists, and we denote by *c*_1_ the minimal wave speed. Similarly, if 0 *< Q < a*_2_, *c*_2_ is the minimal wave speed for the second equation.

Consider the solution *u*(*x, t*) of the Cauchy problem for equation (4.19) with a sufficiently large initial condition *u*(*x*, 0) vanishing at infinity, *u*(*x*, 0) → 0 as *x* ± ∞. Then this solution approaches two waves propagating in the opposite directions and *u*(*x, t*) → *u*_2_ as *t* → ∞ uniformly on every bounded interval. Therefore, *J*_1_(*u*) ∼ 2*u*_2_*c*_1_*t* as *t* → ∞. Similarly for equation (4.20), *J*_2_(*v*) ∼ 2*v*_2_*c*_2_*t* as *t→ ∞*. We call such solutions expanding wave solutions.

We have

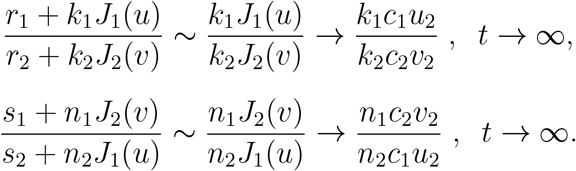

System (4.2), (4.3) can be reduced asymptotically for large time to system (4.19), (4.20) if the following relations are satisfied:

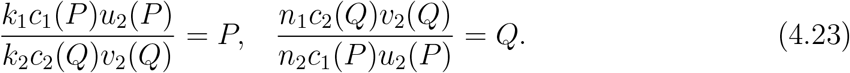

We take into account here that *c*_1_ and *u*_2_ depend on *P, c*_2_ and *v*_2_ depend on *Q*, Multiplying these two equalities, we obtain

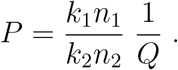

Since *Q* ∈ (0, *q*_∗_), then

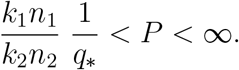

On the other hand, *P* is defined in the interval (0, *p*_∗_). These two intervals intersect if

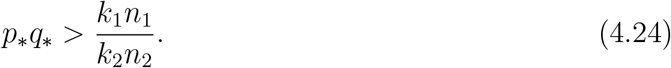

It is the same condition as in Proposition 4.4 for the existence of pulses. Assuming that it is satisfied, we consider the equation

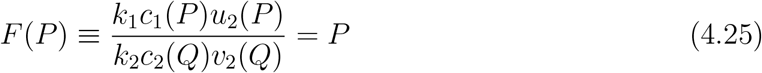

with respect to *P*, where

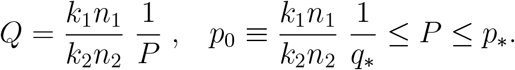

Note that *c*_1_(*p*_0_) *>* 0, *c*_1_(*p*_∗_) = 0. Moreover, if *P* = *p*_0_, then *Q* = *q*∗, and *c*(*Q*) = 0. If *P* = *p*_∗_, then *Q < q*_∗_, and *c*(*Q*) *>* 0. Therefore *F* (*p*_0_) = ∞, *F* (*p*_∗_) = 0. Moreover, functions *u*_2_(*P*) and *v*_2_(*Q*) are bounded from above and from below by some positive constants. Therefore, equation (4.25) has a solution.

#### Theorem 4.6

*If system of equations (4.2), (4.3) has an expanding wave solution, then inequality (4*.*24) is satisfied*.

This assertion does not give sufficient conditions of the existence of such solutions. However, for constant *P* and *Q* such solutions do exist. Their existence can be proved for time-dependent *P* and *Q* converging to some limits. So, we can expect existence of such solutions for the coupled problem (4.2), (4.3).

Conditions of Theorem 4.5 exclude the existence of such solutions, while conditions of Proposition 4.4 admit them. Combining these results with the results of numerical simulations discussed below, we can conclude that waves with positive speeds (Theorem 4.6) exist together with unstable pulses (provided by Proposition 4.4), but not with stable pulses. This is similar for the scalar equation and monotone systems for which unstable pulses exist if and only if the wave speed is positive [35]. Stable pulses do not exist for the scalar equations and monotone systems [31].

The results on tissue growth can be summarized in terms of the non-dimensional parameter

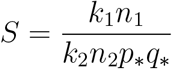

characterizing the strength of the feedback. If *S >* 1, then the solutions converge to a stable pulse. If the inequality is opposite, they propagate as a wave.

### 4.3 Numerical simulations

According to the analytical results presented above, inequality

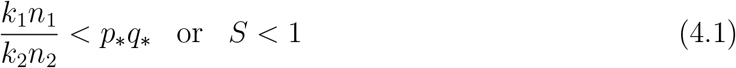

is associated with the existence of expanding solutions, that is, waves with positive speeds. Theorem 4.6 proves that it is a necessary condition of the existence of such solutions. Numerical simulations confirm their existence (Figure 8). Note that the wave speeds for the two components of solutions are, in general, different from each other.

**Figure 8:**
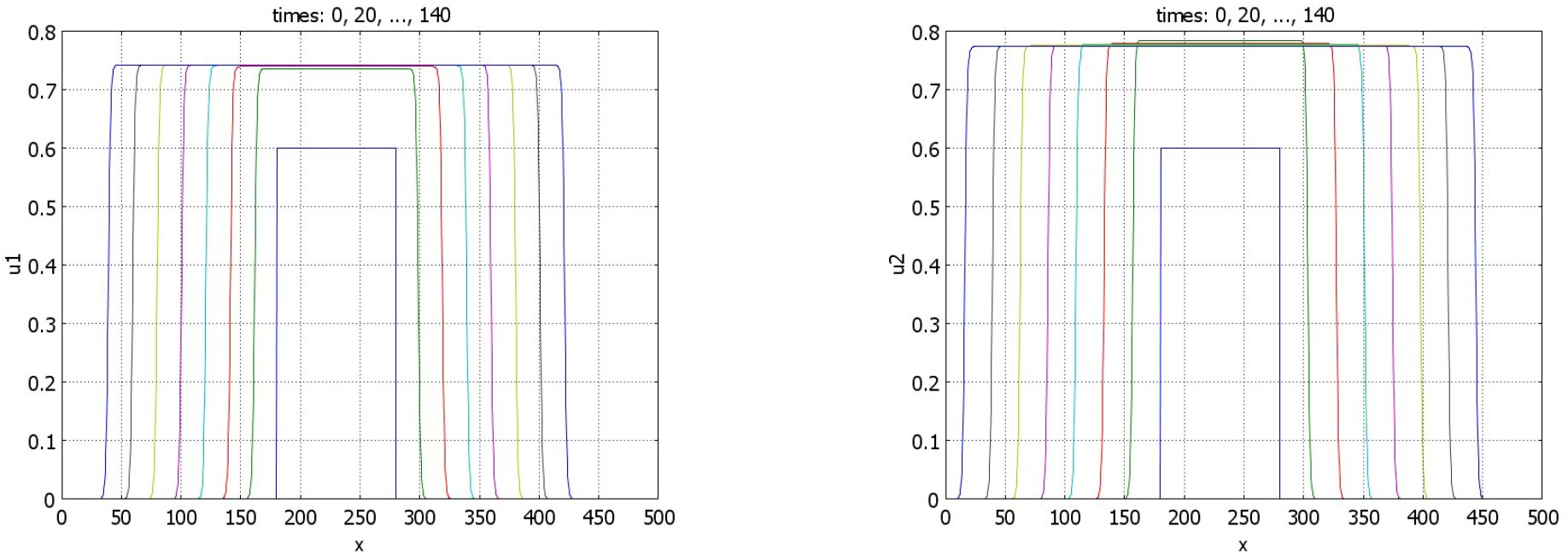
Time profiles of the function *u*_1_ (left) and of the function *u*_2_ (right) for *r*_1_ = *s*_1_ = 0.5, *r*_2_ = *s*_2_ = 1.5, *k*_1_ = 1.2, *n*_1_ = 0.4. The initial condition is a piece-wise constant function; *a*_1_ = *a*_2_ = 1, *b*_1_ = *b*_2_ = 4, *k*_2_ = 1, *n*_2_ = 0.5. For these values of parameters *p*_∗_ = *q*_∗_ = 1.5 and *S* ≈ 0.427. Dynamics of solutions corresponds to expanding waves.

If inequality (4.1) is opposite and conditions of Theorem 3.5 are satisfied, then there exist stationary pulses (Figure 9). We note that the components *u*_1_ and *u*_2_ of solution of problem (4.4)-(4.6) are coupled only through the values of their integrals. Therefore, for any solution *u*_1_(*x*), *u*_2_(*x*) of this problem, then functions *u*_1_(*x*), *u*_2_(*x* + *h*) also satisfy it for any real *h*. This property is illustrated in Figure 9 (left). Increase of the value of *k*_1_ decreases the component *u*_1_ of the solution (Figure 9, right). It also leads to the decrease of *u*_2_ through the integral *J*(*u*_1_).

**Figure 9:**
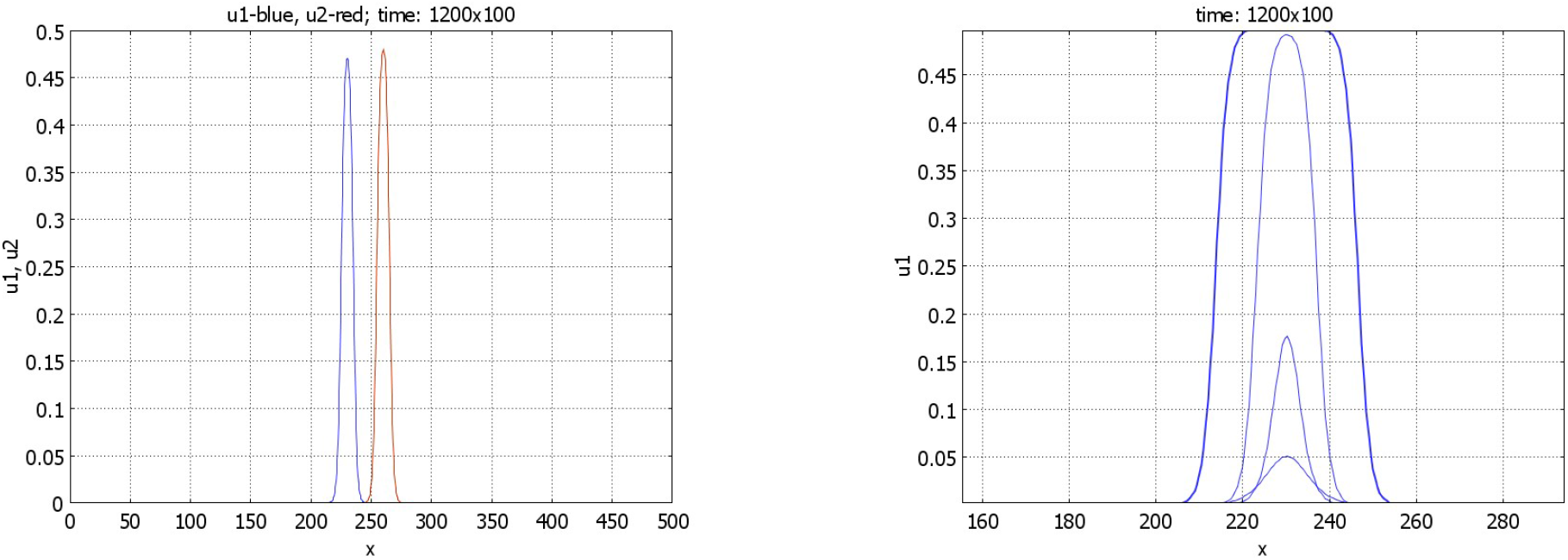
Left panel: profiles of the pulses *u*_1_ and *u*_2_ for *r*_1_ = *s*_1_ = 0.5, *r*_2_ = *s*_2_ = 1.5, *k*_1_ = 2, *n*_1_ = 1. The initial data for *u*_1_ and *u*_2_ are given by rectangles shifted by 30 space units. Right panel: profiles of the *u*_1_ pulses for *r*_1_ = *s*_1_ = 0.5, *r*_2_ = *s*_2_ = 1.5, and fixed *n*_1_ = 2. The maximal value of the profiles decreases with *k*_1_ equal respectively to 0.75, 1, 2, 3. Other parameters: *k*_2_ = 1, *n*_2_ = 0.5, and *S* ≈ 1.78.

Let us now consider the case *r*_1_ = *s*_1_ = *r*_2_ = *s*_2_ = 0 (cf. Proposition 4.4). If condition (4.1) is satisfied, then numerical simulations show that the stationary pulses are unstable, and we observe wave propagation (Figure 10). If inequality (4.1) is opposite, waves with negative speeds are observed (Figure 11). In agreement with Proposition 4.4, there are no stationary pulses.

**Figure 10:**
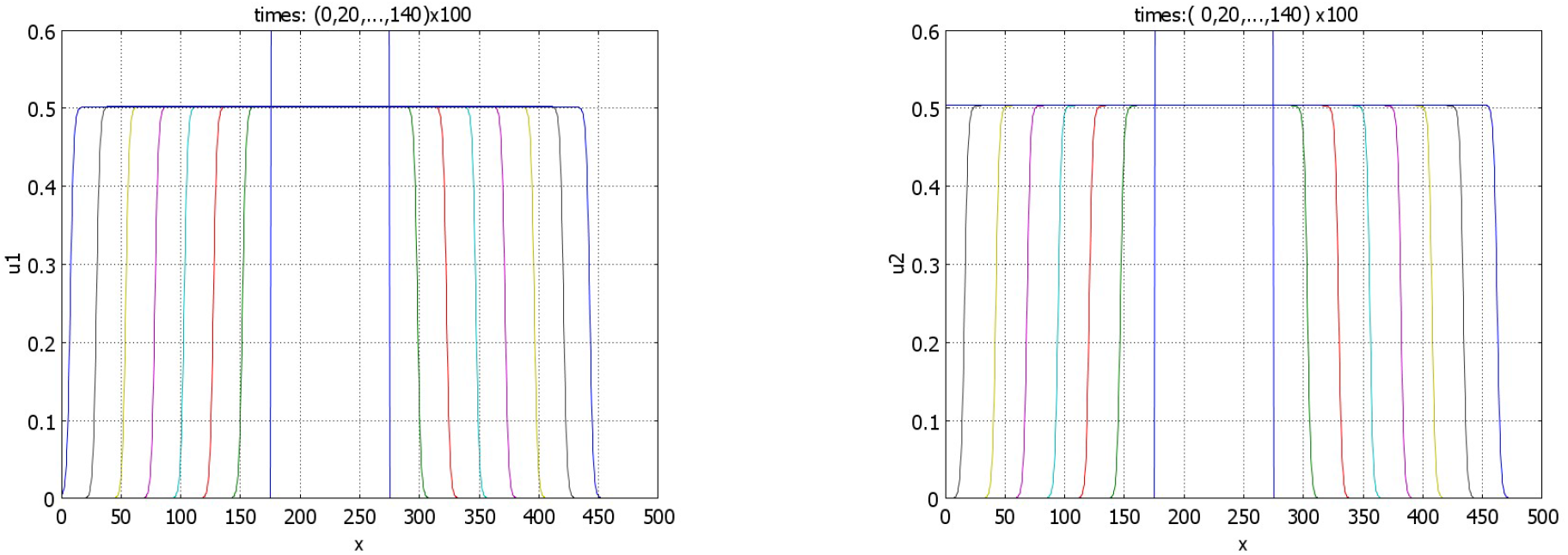
Time profiles of the function *u*_1_ (left) and of the function *u*_2_ (right) for *r*_1_ = *s*_1_ = *r*_2_ = *s*_2_ = 0, *k*_1_ = 1.6, *n*_1_ = 0.7, *k*_2_ = 1, *n*_2_ = 0.5. The initial conditions are given by piece-wise constant functions, *S* ≈ 0.996.

**Figure 11:**
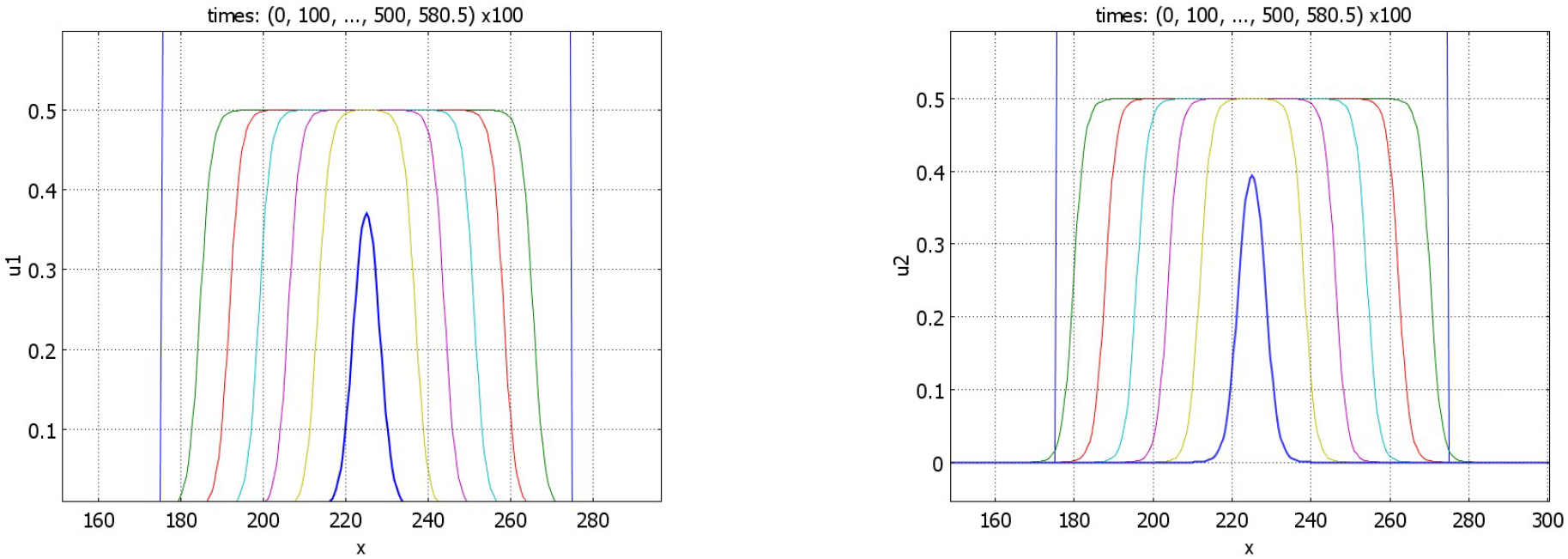
Time profiles of the function *u*_1_ (left) and of the function *u*_2_ (right) for *r*_1_ = *s*_1_ = *r*_2_ = *s*_2_ = 0, *k*_1_ = 1.67725, *n*_1_ = 0.6709. The initial condition is a piece-wise constant function. At time *t* = 581 100 the functions *u*_1_(*x, t*) and *u*_2_(*x, t*) are point-wise smaller than 10^−5^: the solution converges to zero. The values of parameters: *a*_1_ = *a*_2_ = 1, *b*_1_ = *b*_2_ = 4, *k*_2_ = 1, *n*_2_ = 0.5, *p*_∗_ = *q*_∗_ = 1.5, and *S ≈* 1.00023. In agreement with Proposition 4.4, no pulse solution is observed.

Convergence of solution to a wide flat pulse is shown in Figure 12. In the beginning of simulation, solution resembles a travelling wave. After some time, it slows down and stops. Biologically, this example illustrates coordinated growth of two tissues to their final sizes. We use this example for numerical simulations of tissue regeneration. We cut the first tissue, while the second tissue is not changed. In the model, this means that we consider the initial condition with a partially truncated first component, while the initial condition for the second component represents the same stationary pulse as in the previous simulation. The solution of this problem converges to the same pulse solution. The integrals of the first and second components of the solution are shown in Figure 13. The integral of the first component monotonically converges to its stationary value, while the integral of the second component decreases in the beginning and grows later. This decrease in the beginning of simulations shows that the second tissue decreases its size to adjust to the first tissue.

**Figure 12:**
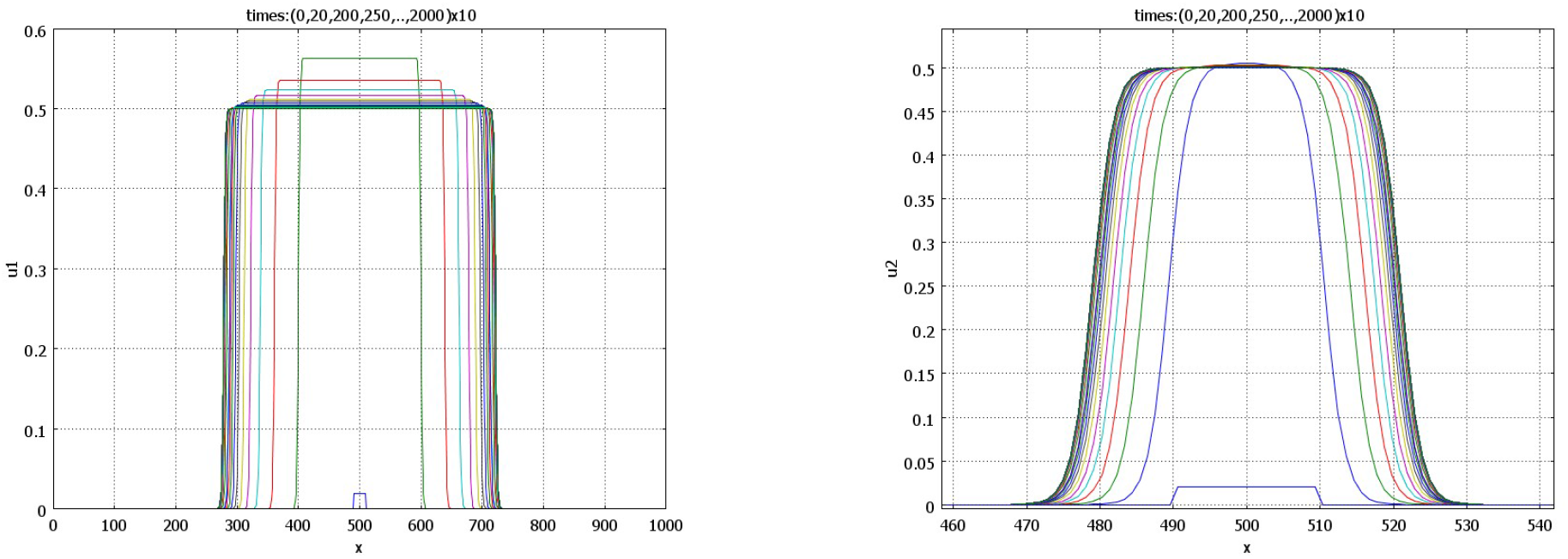
Time profiles of the function *u*_1_ (left) and of the function *u*_2_ (right) for *r*_1_ = 0.5, *s*_1_ = 0.5, *r*_2_ = 1.5, *s*_2_ = 1.5, *k*_1_ = 0.15, *n*_1_ = 8. The initial condition is a piece-wise constant function. Solution converges to the stable pulse. Note that the *x*-scales in the two figures are different. The values of parameters: *a*_1_ = *a*_2_ = 1, *b*_1_ = *b*_2_ = 4, *k*_2_ = 1, *n*_2_ = 0.5, *p*_∗_ = *q*_∗_ = 1.5, and *S* ≈ 1.067.

**Figure 13:**
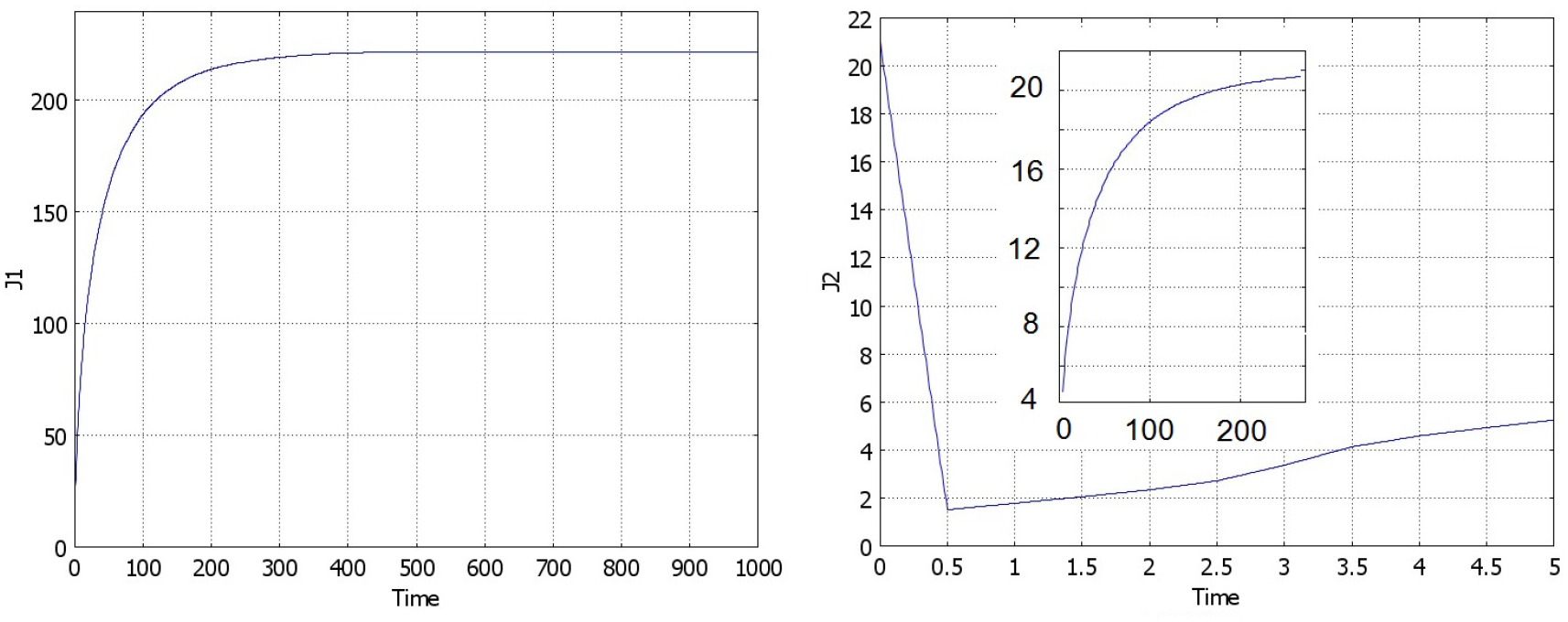
Integrals *J*(*u*_1_) (left) and *J*(*u*_2_) (right) in numerical simulations of tissue regeneration. For the same values of parameters as in Figure 12, the initial condition *u*_1_(*x*, 0) corresponds to the truncated stationary pulse, and for *u*_2_(*x*, 0) to the complete pulse. The solution converges to the same stationary pulse. The integral of the first component of the solution monotonically increases, while for the second component, it first decreases, then grows to its stationary value.

## 5 Discussion

As indicated above in the introduction, tissue growth control can be influenced by tissue cross-talk and negative feedback. We will discuss here the biological mechanisms of this feedback and their realization in the model.

### 5.1 Chalones, growth-inhibitory feedback mechanisms, functional feedbacks

Chalones are tissue-specific, secreted factors that inhibit further cell proliferation in the tissue of origin. The concept of chalones originated with the hypothesis that organ size is regulated by locally produced substances that act to limit growth when a critical mass is achieved [32, 38]. Chalones are secreted in proportion to the size or cell number of the tissue.

As the tissue grows, their concentration increases locally or systemically, inhibiting further proliferation.

Chalones are critical for maintaining tissue homeostasis and preventing excessive growth. The disruption of chalone signaling can result in unregulated tissue growth, contributing to tumorigenesis. For example, loss of myostatin expression in some cancers correlates with tumor progression and metastasis.

The concept of chalones is being discussed during already more than a century with some examples and counter-examples (see the reviews in [2, 38, 39]). In the modern biological literature, it is accepted that it can work for some organs (see Table 1 in [32]), but it is not universal. Other mechanisms are also involved in tissue growth regulation depending on the tissue in question and physiological conditions (embryogenesis, development, trauma, neoplasia).

Growth is often regulated through negative feedback loops, which maintain a balance between proliferation and tissue size. These feedback loops can involve systemic signals (e.g., hormones) or local factors (e.g., chalones, mechanical stress).

Local feedback mechanisms include contact inhibition, where high cell density suppresses cell division, is a classic feedback mechanism. Mechanical cues, such as stiffness of the extracellular matrix (ECM), can signal cells to reduce proliferation as tissue tension increases. Molecules like FGF (fibroblast growth factors) and VEGF (vascular endothelial growth factor) operate in local feedback loops to regulate organ development and maintain proportions. Parabiosis studies in mice (joining circulatory systems of two animals) have shown that systemic factors can rejuvenate aged tissues, but intrinsic local feedback mechanisms primarily govern growth control.

Another growth control is based on functional feedback mechanisms that use organ performance or output to regulate growth. Unlike chalones, which depend on mass or cell number, functional feedback ensures that tissue size matches physiological needs.

Examples of functional control include liver and bile acid flux. Bile acids synthesized by the liver are recirculated through the enterohepatic system. When bile acid flux increases due to enhanced digestion demands, hepatocyte proliferation is induced to expand liver size. Conversely, reduced bile acid flux signals that the liver has reached a sufficient size.

Another example concerns kidney and functional compensation. Following unilateral nephrectomy, the remaining kidney undergoes hypertrophy to compensate for lost function. The signal for this compensation may involve increased serum creatinine levels or local mechanical stress. Importantly, this compensatory growth primarily involves hypertrophy rather than hyperplasia. In thyroid and endocrine glands, negative feedback loops involving hormone levels (e.g., TSH and thyroid hormone) regulate the size and activity of endocrine organs. Hypothyroidism leads to compensatory thyroid hypertrophy (goiter), while hyperthyroidism suppresses TSH secretion, reducing thyroid size.

In distinction from chalones, functional feedback is performance-driven, whereas chalone mechanisms are based on physical size or cell number. Functional feedback often operates through systemic signaling, as seen in liver regeneration involving bile acids or kidney compensation mediated by metabolic demands.

### 5.2 Models of growth with negative feedbacks

The mechanisms of tissue growth control presented above act through negative feedback. Biological mechanisms of this feedback and their implementations in the models can be different. A common feature of these mechanisms, including functional feedback, is that there is some characterization of the tissue (signal, function) proportional to its size *J*(*u*). If we denote its level by *B*, then we obtain equation (2.2) for its time evolution.

From this point on, these mechanisms begin to differentiate. Global (systemic) feedback can be expressed by another signaling molecule *C* produced by some other organ and described by equation (2.3). This endocrine signaling can up-regulate cell death in equation (2.1) or down-regulate its proliferation.

Functional feedback, which is also systemic, is determined by some given value *B*_0_ required by the organism. Therefore, tissue growth is proportional to (*B*_0_ − *B*) or, in the dimensionless form, to (1 − *J*(*u*)) entering as a factor in the tissue proliferation rate. From the modelling point of view, this is similar to systemic feedback down-regulating cell proliferation in the previous paragraph.

Local (autocrine, paracrine) feedback, associated with chalones (though their action can also be systemic), decreases cell proliferation. However, the quantity of substance *B* with respect to a single tissue cell, that is *B/J*(*u*) remains constant since *B ∼ J*(*u*). Therefore, its action does not depend on the tissue volume, and cannot control its growth.

There is a principal difference between the local feedback, where the *total* quantity of substance *B* is proportional to *J*(*u*), and systemic feedback, where the *concentration* (level) of *C* in blood (or in the whole organism) is proportional to *J*(*u*). In the first case, as discussed above, the quantity of *B* with respect to a single cell remains approximately constant, and feedback intensity does not depend on the tissue volume (Figure 14). But relative concentration of *B* can increase with time and act as growth inhibitor or growth promoter that decreases expression of their growth [32]. In the second case, depletion of *C* in the tissue is negligible compared to the whole organism, we do not divide *C* by *J*(*u*). In this case, feedback intensity depends on the tissue volume *J*(*u*).

**Figure 14:**
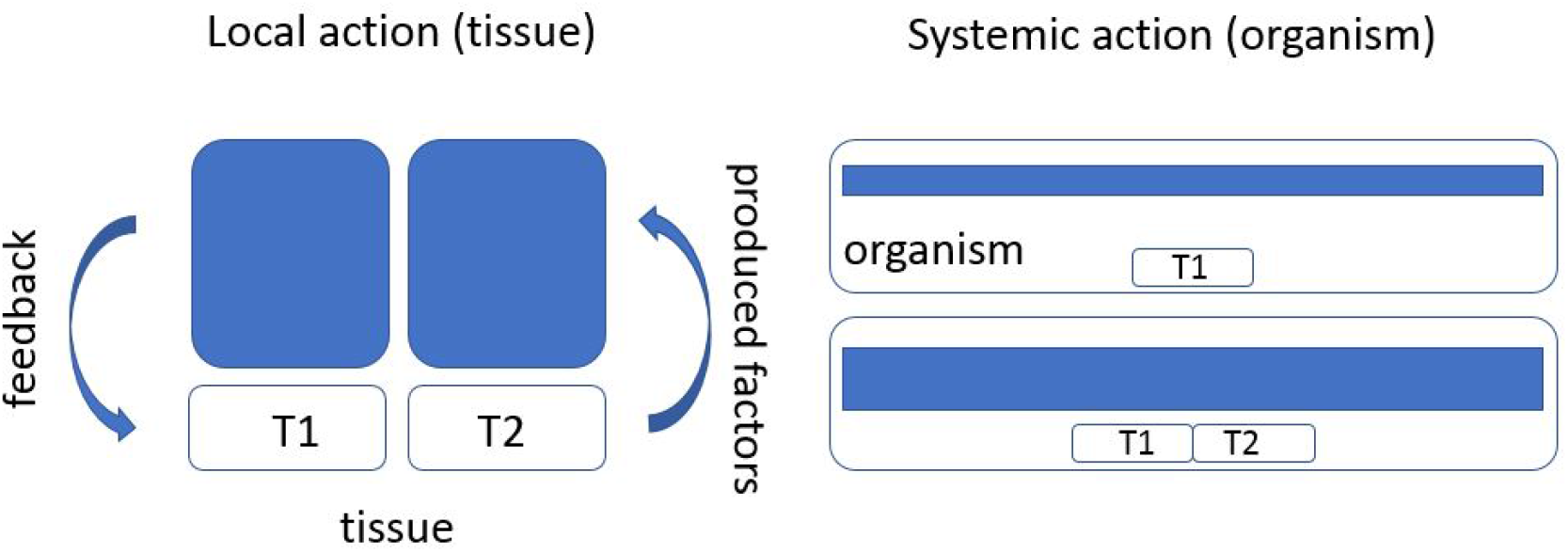
Local (left) and systemic (right) feedback on tissue growth by some factors produced by the tissue. In the local case, increasing the tissue twice (*T* 1 + *T* 2), increases the total amount of the produced factor. However, its amount with respect to the unit tissue volume remains the same. Therefore, local feedback cannot determine the tissue size. In systemic feedback, instead of the total amount of the produced factor in the whole organism, we measure its level (concentration). Increasing twice the tissue, we also increase twice this level. Its feedback on the unit tissue volume also increases.

Thus, we model systemic feedback, the mechanism of which can be related to chalones and to other negative feedbacks, or to the functional feedback.

### 5.3 Modelling results and biological interpretations

#### 5.3.1 Single tissue with systemic feedback

##### Spatially-distributed solutions and tissue differentiation

The question of tissue differentiation in a growing embryo was first addressed by A. Turing in his seminal work [33]. He proposed a mechanism based on diffusion-driven instability, which arises from the interplay between long-range inhibition and short-range activation. This mechanism results in the formation of spatially periodic patterns. In spite of the enormous interest to Turing structures in mathematical biology, the mechanisms of cell differentiation in a growing embryo are likely to be different [41].

In Section 3.1, we proposed an alternative mechanism for the emergence of spatial structures, which bifurcate from a spatially homogeneous solution. Biologically, this mechanism relies on local cell communication, which promotes cell proliferation, and global negative feedback, which stimulates cell death.

The global feedback operates through the integral term *J*(*u*), which modifies the eigen-value distribution of the linearized problem. Specifically, it decreases the eigenvalue *λ*_0_ with the largest real part, as the corresponding eigenfunction *ϕ*_0_ is a positive constant, and *J*(*ϕ*_0_) *>* 0. However, for all other eigenfunctions, *ϕ*_*i*_(*x*) = cos(*πnx/L*), the integral vanishes, leaving the eigenvalues unchanged. Consequently, under certain parameter conditions, the second eigenvalue *λ*_1_ surpasses *λ*_0_. This instability in the spatially homogeneous solution then leads to the formation of spatial structures.

A more detailed comparison between this instability and Turing instability is provided in [30], particularly regarding its dependence on interval length and the rates of cell division and death.

It is worth noting that in equation (2.1), *u*(*x, t*) is interpreted as cell concentration, with the diffusion term representing random cell motion. Alternatively, this variable can also be understood as the concentration of an autocrine signaling molecule or other locally produced molecules that regulate cell division or differentiation.

##### Existence of pulses

Spatially distributed solutions bifurcating from a constant solution take the form of pulses further into the instability region. This bifurcation occurs in problems defined on a bounded interval but not on the whole axis. On the whole axis, the integral of a positive constant solution is not defined, and for the zero solution, bifurcation does not occur. Therefore, in this context, we focus on the existence of pulses rather than their bifurcation.

The existence of pulses depends on the properties of the feedback function *g*(*J*(*u*)). Under the conditions specified in Theorem 3.2, equation (3.12) admits a solution, which ensures the existence of a pulse. However, the uniqueness of this solution is not guaranteed by the theorem and may not generally hold. In the generic case, where solutions do not overlap, their number is odd. If the conditions on the feedback function are not met, the number of solutions becomes even, potentially resulting in no solutions at all. Notably, two pulse solutions were identified in [34] for a different function *F* in a population dynamics model.

One of the key conditions for the existence of pulses is *b > a*. Biologically, this implies that the effect of local cell-cell communication on the proliferation rate must be sufficiently strong. This phenomenon has some resemblance to the emergence of Turing structures, where short-range activation and long-range inhibition drive pattern formation. In this case, the long-range inhibition is provided by the negative feedback.

##### Stability of pulses

If *g*(*J*(*u*)) ≡ const, that is, equation (2.1) does not depend on the integral, then the pulse solution is unstable. Indeed, the eigenfunction of the zero eigenvalue of the corresponding eigenvalue problem is the derivative of the pulse solution. Therefore, it has variable sign. On the other hand, the eigenfunction corresponding to the eigenvalue with the maximal real part is positive [31]. Hence, *λ* = 0 is not the principal eigenvalue and, consequently, there is a positive eigenvalue. This means that the pulse solution is unstable. Introduction of the integral term in the equation can make this solution stable (see also [36]). This is not proved mathematically but confirmed in numerical simulations. Stability of pulses can be related to existence and bifurcation of solutions of equation (3.12).

Convergence to wide flat pulses is appropriate for tissue growth control. The solution dynamics looks like a wave propagation. After some time, its speed decreases and the propagation stops (Figure 6). This growth arrest is determined by the negative feedback through the tissue size *J*(*u*).

It is important to note that such behavior of solution is observed for any small initial condition. Due to the time dependence of the integral *J*(*u*), the nonlinearity in the equation is of the monostable type in the beginning of the simulation and of the bistable type some time later. This change in the type of equation provides growth of solution in the beginning and existence of a stable pulse for large time. From the biological point of view, this is appropriate for the description of tissue growth in embryogenesis, and in modelling of tumor growth.

#### 5.3.2 Coordinated growth of two tissues

All tissues and organs in the growing organism precisely correspond to each other in their sizes and functionality. The question about the mechanisms of this coordination is largely discussed in the biological literature (see the discussion above). In the model, tissue growth regulation is provided by a negative feedback by each tissue on itself, and by a positive feedback on the other one (see [38] for the biological discussion). The former controls tissue convergence to a final size (similar to the model of a single tissue), while the latter determines the proportional growth between the two tissues. The question about other possible models remains open. Note that for a model in population dynamics stable pulses can exist in the case where all interactions are negative [37].

In this work, we considered tissue cross-talk through the rate of cell death. Their interaction in the cell proliferation rate will be studied in the future works.

##### Existence of waves and pulses

Dynamics of solutions in the model of two tissue is determined by the parameter *S* which characterizes the feedback in the rate of cell death for both tissues. Larger values of this parameter correspond to stronger feedback. If *S >* 1, then there exists a pulse solution (Theorem 4.5). Numerical simulations show that such solutions are stable. Thus, in the case of strong feedback, the final tissue sizes are finite.

Weak feedback in cell death corresponds to wave propagation (Theorem 4.6). In this case, existence of pulses is not observed in numerical simulations. We can conclude that weak feedback leads to the unlimited tissue growth.

Additional condition of the existence of pulses in Theorem 4.5, *r*_1_*/r*_2_ *< a*_1_, *s*_1_*/s*_2_ *< a*_2_, signify that in the beginning of tissue growth, when the feedback is negligible, the cell proliferation rate exceeds the death rate. Therefore, tissue growth can occur with any small initial condition. This corresponds to the biological understanding of this process.

An interesting case is provided by the conditions *r*_1_ = *r*_2_ = *s*_1_ = *s*_2_ = 0 which cannot be considered as a particular example of the more general case discussed above. This case is different from the point of view of the imposed conditions and of the corresponding results. The pulses can exist in this case but they are unstable, and wave propagation is observed in this case in numerical simulations.

##### Tissue growth rate

Tissue growth dynamics exhibit distinct phases: an initial exponential growth, a phase of constant growth, and a final phase of deceleration as the tissue stabilizes to its ultimate size [32, 38]. These phases reflect both biological and model-based representations of tissue growth (Figures 5, 6, 13). Early exponential growth is driven by rapid cell proliferation, supported by abundant resources and minimal constraints. As cell density increases, however, growth transitions to a constant rate due to density-dependent inhibition of cell division. This phenomenon reflects contact inhibition and competition for limited resources, which collectively cap the proliferation rate. In the final phase, systemic feedback mechanisms come into play, curbing growth through the promotion of cell death or senescence. Negative feedback from systemic factors, such as signaling molecules or hormones, ensures that tissues do not exceed their optimal size, maintaining homeostasis.

In the modelling, the transition from the exponential growth to a constant growth rate occurs due to the logistic term in the proliferation rate. Instead of simultaneous cell division in the whole volume at the first stage, we observe predominant cell division at the exterior part of the tissue and lateral growth lile a reaction-diffusion wave. At the third stage, this propagation slows down and stops due to the integral term describing negative systemic feedback.

In regeneration, tissue growth dynamics involve additional complexities. When one tissue is ablated, compensatory mechanisms in the surrounding tissues are activated. These mechanisms often result in an initial decrease in the size of adjacent tissues, enabling proper recalibration and eventual size restoration [40] (cf. Figure 13). This decrease is thought to be regulated by systemic feedback that synchronizes growth rates across tissues, ensuring proportional regeneration. The interplay of local cellular factors and systemic regulatory signals highlights the intricate control mechanisms underlying tissue growth and regeneration, providing insights into both normal development and potential therapeutic interventions. Understanding these processes is essential for unraveling the principles governing tissue homeostasis and recovery after injury.

## 6 Conclusions

This study presents a comprehensive mathematical framework for understanding tissue growth regulation via endocrine signaling. Through the development of reaction-diffusion models, we examined the interplay between local and systemic feedback mechanisms in controlling cell proliferation and tissue size. The models elucidate several critical phenomena, including:

Emergence of spatial structures. The analysis reveals that tissue growth and differentiation can be driven by local cell-cell communication, coupled with global feedback mechanisms. This combination leads to stationary pulse solutions, offering a novel explanation for tissue differentiation beyond classical Turing instability.

Regulation through negative feedback. The results highlight the importance of systemic feedback mechanisms, such as endocrine signaling, in stabilizing tissue growth and achieving proportionate sizes. Stability and existence of pulse solutions depend on the strength and type of feedback, with strong feedback ensuring finite tissue sizes, while weaker feedback leads to unlimited growth.

Coordinated growth of tissues. The study demonstrates that inter-tissue signaling can coordinate the growth of multiple tissues, ensuring harmonious development. Positive feedback between tissues amplifies this coordination, while negative feedback determines the final tissue size.

Bifurcation and stability of solutions. The existence and stability of stationary pulses and traveling wave solutions were characterized under various parameter regimes. These findings provide a mathematical basis for understanding phenomena such as growth arrest, regeneration, and tumor expansion.

Biological implications. The models suggested in this work offer insights into diverse biological processes, including embryogenesis, organ size regulation, and pathological conditions such as cancer. This framework also underscores the role of tissue cross-talk in maintaining systemic homeostasis.

Future work will extend these models to include additional biological complexities, such as functional feedback mechanisms and specific signaling pathways, to further enhance our understanding of growth regulation and tissue coordination.

## Acknowledgments

The last author has been supported by the RUDN University Strategic Academic Leadership Program.

